# Cancer-driven neutrophil priming couples systemic epithelial regenerative programs with pre-metastatic niche formation

**DOI:** 10.1101/2025.10.20.683483

**Authors:** Nicolas Rabas, Victoria L. Bridgeman, Anisha Ramessur, Constandina Pospori, Marc Hennequart, Felipe S. Rodrigues, Ana Farias, Rute M. M. Ferreira, Konstantinos Axarlis, Olamide Nduka, Probir Chakravarty, Nathalie Legrave, Izadora Furlani, Raoul Charles Coombes, James MacRae, Ilaria Malanchi

**Affiliations:** Tumour-Host Interaction Laboratory; Bone Marrow Dynamics Laboratory; Tumour and Host Metabolism Laboratory; Bioinformatics Core; Metabolomics, The Francis Crick Institute, London, United Kingdom; Respiratory Infections Section, National Heart and Lung Institute; Department of Surgery and Cancer, Imperial College London, London

## Abstract

Cancer progression involves systemic changes that extend beyond the primary tumour. Through cancer-induced systemic conditioning, breast tumours generate subclinical alterations in distant organs that facilitate metastatic seeding and pre-metastatic niche formation. Neutrophils, mobilized through cancer-driven emergency granulopoiesis, actively contribute to this process. In this study we extend the concept of neutrophil-dependent conditioning beyond pre-metastatic sites, uncovering a broader systemic regenerative activation that links inflammation, tissue regeneration, and metastasis. This activation manifests as enhanced epithelial progenitor activity, measured by increased organoid formation, across multiple organs, including those with low risk of breast cancer metastasis. This neutrophil-dependent perturbation in lung alveolar progenitors and intestinal epithelial lineage commitment, is an indication of an altered organ physiology, enhancing tissue resilience to injury. Moreover, we identify UPP1 expression, which exclusively characterizes neutrophils generated through emergency granulopoiesis, as a key factor sustaining high translational activity in neutrophil progenitors and enabling the full acquisition of cancer-primed properties. Consequently, neutrophil loss of UPP1 reduces both their lung pro-metastatic function and their capacity to activate alveolar progenitors. Mechanistically, this involves interactions between cancer-primed neutrophils and platelets, which localize within lung interstitial spaces near alveolar cells to stimulate epithelial progenitor activity.

Together, these findings uncover a previously unrecognized tumour-induced systemic conditioning in which neutrophils coordinate epithelial regenerative activation as part of a pro-metastatic epithelial niche, with UPP1 as a key determinant of their cancer-primed state.

## Introduction

Tumours release a broad repertoire of systemic factors that perturb host physiology far beyond the primary site^1^. Through direct release of soluble factors and extracellular vesicles, or indirectly through perturbation in immune cell production and activity, primary tumours induce subclinical changes in distant organs that increase the chance of future metastatic colonisation and therefore generate pre-metastatic niches^1,2^. These niches are characterised by coordinated alterations in vascular permeability, stromal activation, extracellular matrix remodelling, immunosuppressive microenvironment and metabolite supply, which collectively condition organs for subsequent metastatic colonisation^1^. While the molecular mechanisms underlying niche formation are increasingly defined, the underlying physiology of the conditioned state remains underexplored.

Many hallmark features of the pre-metastatic niche mirror those that accompany regenerative responses after injury. For example, fibronectin and Tenascin-C deposition^3,4^ and lysyl oxidase-mediate collagen crosslinking^5^ are classical extracellular matrix changes shared between the pre-metastatic lung and repairing tissues^6^. Also the well-known pro-tumorigenic activity of neutrophils in cancer can be viewed in parallel with their role in wound healing^7,8^. Therefore, the cancer mediated pre-metastatic conditioning resemble a state of chronic, low-grade injury response that inadvertently creates conditions favourable for metastatic colonisation. Indeed, metastatic cells needs this wound-healing-like response and locally further induce stromal activation, angiogenesis, ECM remodelling, and immune modulation^9^. Within this context, epithelial regenerative program can be detected early in metastatic niche establishment^10^, which reprogram epithelial cells to reacquire multilineage differentiation potential, reinforcing the idea that echoing regenerative programs is require for cancer progression^11–13^. While these parallels suggest that a regenerative-like program within distant pre-metastatic niches may underlie cancer-induced distant conditioning, whether this translates into a measurable increase in regenerative activity remains unknown.

Neutrophils are key effectors of tumour-induced systemic conditioning, where cancer-driven perturbation of granulopoiesis leads to the production and mobilisation of cancer-primed neutrophils with long known function in pre-metastatic niche formation^7,14,15^. Beyond this, neutrophils can activate epithelial progenitors during tissue repair, as demonstrated following irradiation-induced lung injury to facilitate metastatic colonisation^16^. Here we tested whether cancer-mobilised neutrophils engage regenerative activation when establishing the pre-metastatic niche.

## Results

### Breast cancer enhances homeostatic epithelial progenitors’ activation of distant organs

To assess whether primary tumour influence the steady-state regenerative potential of distal epithelia, we employed organoid formation assay. In this assay, the number of organoids formed directly reflects the number of epithelial cells within the organ retaining the stemness ability to recapitulate an organ-like structure *ex vivo* at the time of isolation. When comparing cells isolated from different organs from either naïve or MMTV-PyMT tumour-bearing animals at the pre-metastatic stage, we found a consistent increase in organoid-forming efficiency (Figure 1a-e and Extended Data Figure 1a-d). Notably, this phenomenon was not restricted to the organ targeted for spontaneous metastasis, but extended to other tissues, even if rarely hosting metastatic growth, such as pancreas and intestine. An increase in steady state proliferative activity was also detected in the intestine of tumour-baring (TB) mice (Extended Data Figure 1e). These findings indicate that primary breast tumours alter the steady-state progenitor activity of distal organs that is not specifically targeting pre-metastatic niches.

**Figure 1.**
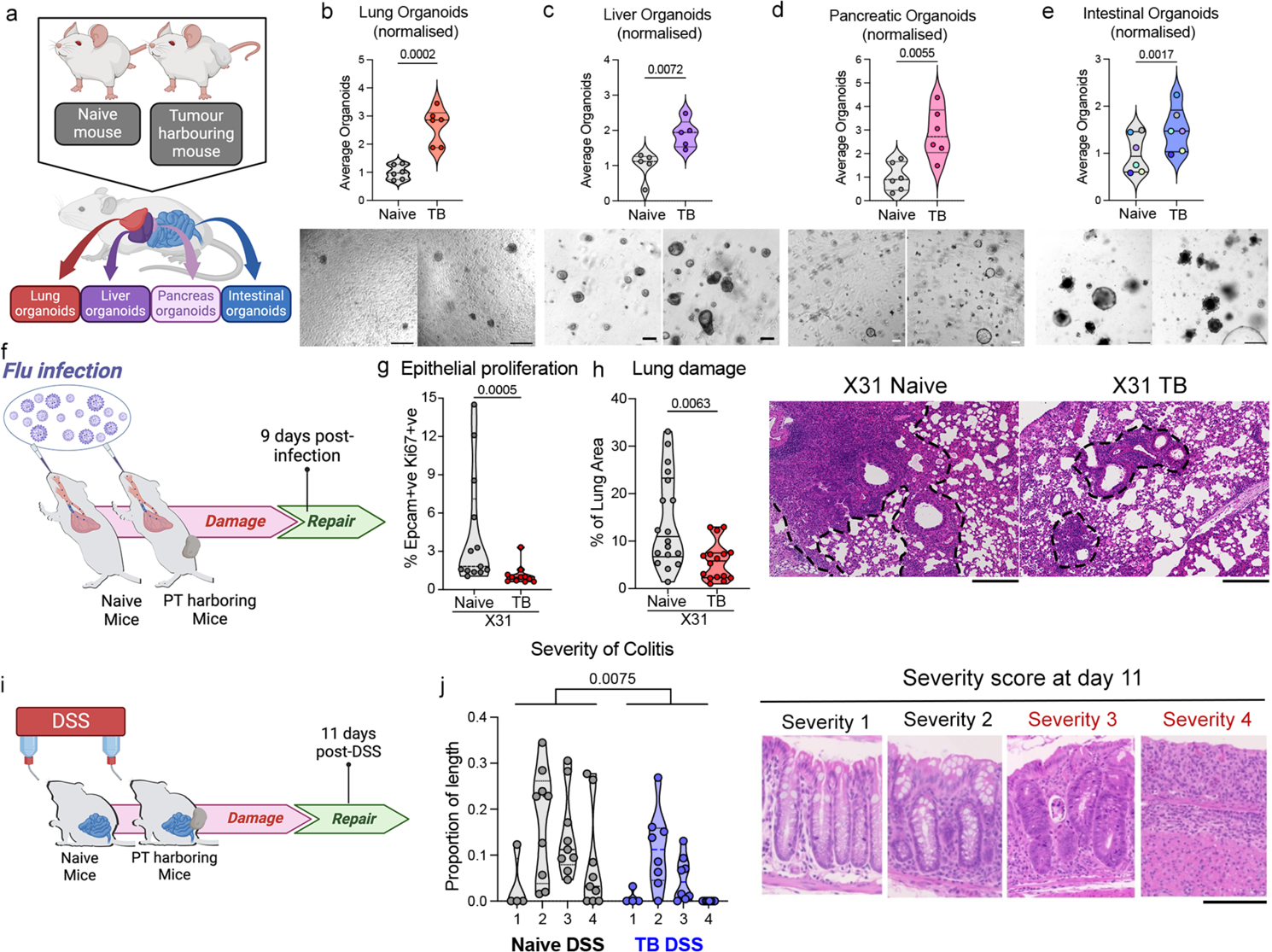
Primary breast tumours induce systemic enhancement of epithelial progenitor activity and tissue resilience to injury. **a)** Schematic representation of multi-organ assessment of organoid formation in naïve and tumour-bearing (PyMT-FVB) mice. **b-e) (Top)** Organoid formation efficiency of cells isolated from lung (b), liver (c), pancreas (d) and intestine (e) of tumour-bearing and Naïve mice. Data are normalized to the average of the control group in each experiment, total organoid numbers shown in Extended Data Figure 1. Violin plots show fold-change in organoid number (n= 6 per group except liver n=5); each dot represents one mouse. For lung, liver, pancreas, student’s t-test was used. Intestinal organoids were analysed with paired t-test (dots coloured by paired experiment). **(Bottom)** Representative bright field images of organoids. **f)** Schematic of influenza infection (X31) with endpoint after 9 days in naïve and tumour-bearing (PyMT C57Bl/6) mice. **g)** Quantification of Ki67+ve epithelial cells in lung during regenerative phase (day 9) following influenza-induced injury (X31) (naïve n=13, tumour-bearing (TB) n=11). Mann-Whitney t-test. **h) (Left)** Histological quantification of the area of lung damage as a percentage of total lung area day 9 (naïve n=18, TB n=16 mice), Mann-Whitney t-test. **(Right)** Representative H&E images dashed lines around regions of lung injury. **i)** Schematic of DSS-induced colitis, naïve and tumour bearing (PyMT-FVB) mice received DSS drinking water for 7 days with damage assessed 4 days after treatment. **j) (Left)** Histological quantification of the proportion of colon length exhibiting indicated damage severity score: 1, mild inflammation; 2 marked immune infiltrate and crypt abnormalities; 3, intense immune infiltration, crypt hyperplasia, cystic structures; 4, complete epithelial loss with intense immune infiltration, (naïve n=9, TB n=8). Ordinary two-way ANOVA. **(Right)** Representative H&E images illustrate each grade.

We next assessed whether this altered epithelial state could be symptomatic of a general perturbation with potential functional consequences for tissues. To assess a shift in physiologic state, we challenged both the pre-metastatic lung and the cancer-perturbed colon with insults causing a transient injury. Lung injury was induced in naïve and tumour-bearing mice by influenza virus (X31) infection (Figure 1f). Both Naïve and tumour-bearing mice experienced similar disease severity, and the infection did not affect tumour growth over this time period (Extended Data Figure 1f and 1g). When assessing inflammatory response to the infection, we detected an increase total immune cells in tumour-bearing mice, with a notable increase in neutrophils persisting 9 days post infection, while the T cell presence was similar (Extended Data Figure 1h-j). Strikingly, when assessing the lung damage caused by this transient insult, a reduction in both overall epithelial proliferation and tissue damaged areas were detected in lungs of tumour-bearing mice (Figure 1g and 1h). These data indicate an increase resilience to injury-mediated influenza infection in pre-metastatic lungs.

To assess intestinal response to injury, naïve and tumour-bearing mice were treated with DSS for one week to induce colitis. Injury severity was assessed four days after DSS withdrawal, during the recovery phase (Day 11 post DSS) (Figure 1i). While the steady state presence of neutrophils in the colon was increased in tumour bearing mice, upon DSS treatment only a trend toward a higher neutrophil’s infiltration was detected (Extended Data Figure 1k and 1l). Nonetheless, histological scoring of colon injury severity revealed that tumour-bearing mice exhibited a significant reduction in tissue damage, particularly in regions of more severe injury (Severity 3 and Severity 4) (Figure 1j and Extended Data Figure 1m). Therefore, similarly to the post-infected lungs, an increase resilience to DSS-induced colitis was observed in the cancer-conditioned colon.

Collectively, these data indicate that breast cancer systemically induces activation of epithelial progenitor activity across multiple organs. This epithelial conditioning occurs not only in pre-metastatic tissues such as the lung but also in organs unlikely to become future metastatic sites, such as the colon. Most importantly, it reflects a broader physiological shift toward increased tissue resilience to transient injury.

### Cancer-primed neutrophils dependent alteration in the epithelial compartment of lung and intestine

Our previous work and that of others have highlighted the key role of neutrophil in the context of the pre-metastatic niche enstablishment^1,14^. We have also shown that neutrophils actively support regenerative pathways in lung epithelial cells in the context of lung injury caused by radiation^16^. We therefore asked whether neutrophil mobilization was linked to epithelial progenitor activation in the context of cancer, focusing on the lung and intestine, two organs with low and high baseline epithelial turnover, respectively. To this, we performed scRNA-seq from naïve mice, tumour-bearing mice, and tumour-bearing mice where neutrophils were depleted for one weeks. In the lung we focused on the alveolar type 2 (AT2) population, the stem cells of the alveolar epithelium^17^ (Figure 2a and Extended Data Figure 2a). We employed PHATE^18^ to generate an embedding that preserves local and global cell-cell similarities to reveal putative lineage relationships, allowing an unbiased resolution of alveolar progenitor subsets within the AT2 cell population. This analysis resolved three trajectories emerging from a common progenitor pool (cluster 4), which was enriched for canonical alveolar progenitor signatures previously described^19–21^ (Figure 2b, 2c and Extended Data Figure 2b). We next calculated condition-associated likelihoods using MELD^22^ to measure transcriptional perturbations across the trajectory. Interestingly, this unbiased analysis identify cluster 4 of the AT2 progenitors as the most profoundly altered in tumour-bearing mice (Figure 2d). Moreover, an analogous MELD analysis comparing tumour-bearing mice with or without neutrophil depletion, showed a similar enrichment of signal in cluster 4 and cells belonging to its proximal lineage, cluster 2 and 3 (Figure 2e). Accordingly, pathways related to cell plasticity were enriched in AT2 cells in tumour-bearing mice under the influence of neutrophils (Extended Data Figure 2c). Collectively, this analysis reveals a neutrophil-dependent transcriptional perturbation targeting alveolar progenitor cells in the context of breast cancer.

**Figure 2.**
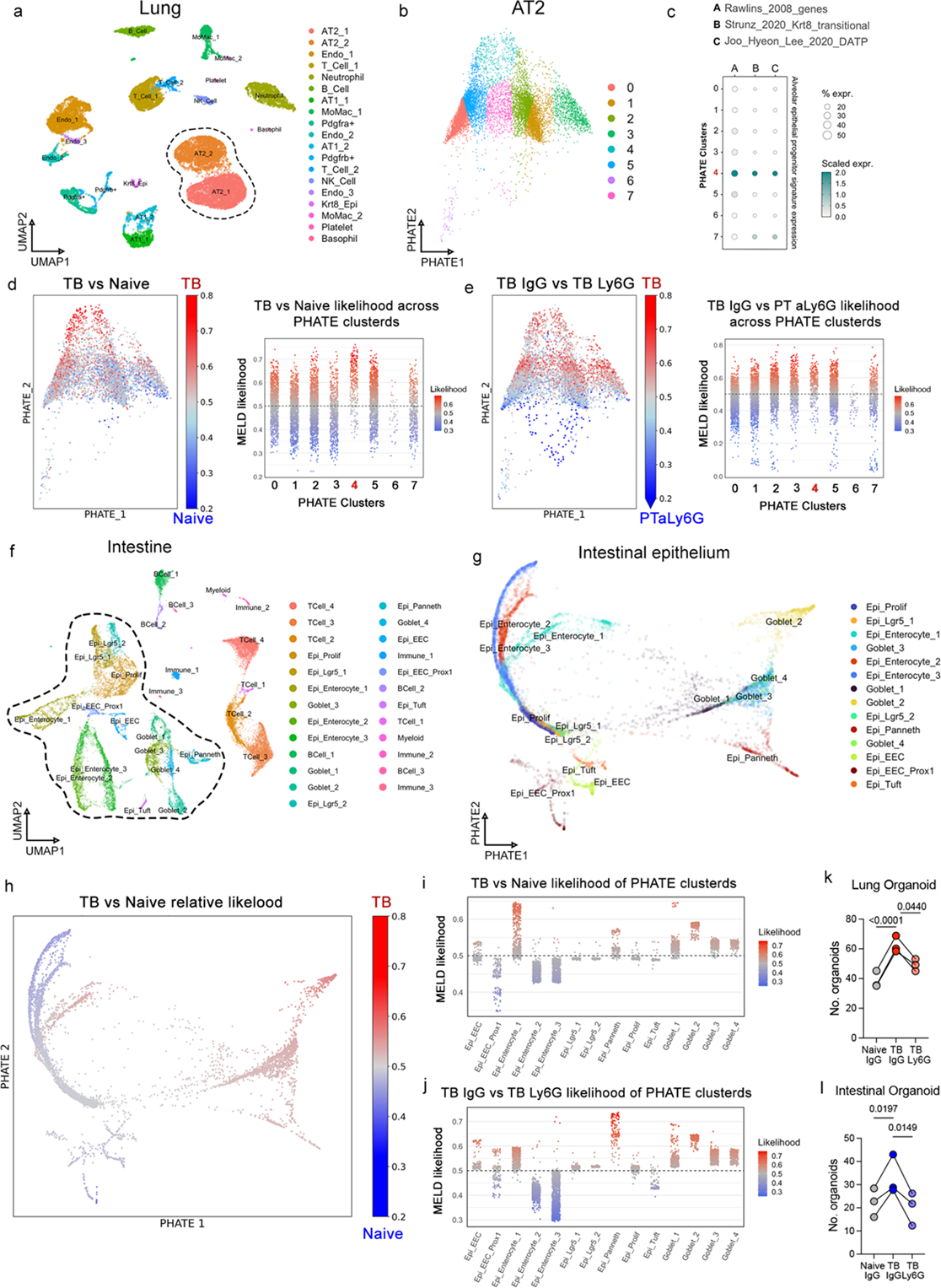
Primary breast tumours induce a neutrophil-dependent perturbation in epithelial compartments of both pre-metastatic lung and intestine. **a)** UMAP embedding of lung scRNAseq data from Naïve IgG, Tumour-bearing (TB) (PyMT-FVB) IgG and TB αLy6G mice. Cells are coloured by cluster with cell type identities indicated in the legend. Alveolar type 2 (AT2) clusters are outlined with dashed line. **b)** PHATE embedding generated from AT2 clusters across Naïve IgG, TB IgG, and TB αLy6G conditions. Cells are coloured by subclusters (0–7). **c)** Expression of alveolar epithelial progenitor-associated signatures across AT2 subclusters shown in (b). Colour indicates scaled expression level. **d) (Left)** MELD analysis showing per cell MELD relative likelihoods represented on the PHATE embedding for Naïve IgG vs TB IgG conditions. Red indicates higher relative likelihood associated with TB IgG condition. **(Right)** Jitterplot shows per-cell TB IgG-associated likelihood across AT2 subclusters. **e) (Left)** MELD analysis comparing TB IgG and TB αLy6G conditions. Red indicates higher likelihood of association with the TB IgG condition. **(Right)** Jitter plot shows per-cell likelihoods across clusters. **f)** UMAP embedding of Intestinal scRNAseq data from Naïve IgG, TB IgG and TB αLy6G conditions. Cells are coloured by cluster with major cell type identities indicated in the legend. Intestinal epithelial clusters are outlined with dashed line. **g)** PHATE embedding generated from epithelial cell clusters across Naïve IgG, TB IgG and TB αLy6G conditions shown in (f), coloured by cell-type identity. **h)** Per-cell MELD relative likelihood represented on PHATE embedding comparing TB IgG and Naïve IgG conditions. Red indicates higher relative likelihood associated with TB IgG. **i-j)** Jitter plots showing per-cell MELD relative likelihoods across intestinal cell-type clusters for (i) TB IgG versus naïve IgG and (j) TB IgG versus TB αLy6G comparisons. Higher likelihood values correspond to TB IgG conditions. **k)** Quantification of organoid formation efficiency of lung epithelial cells isolated from Naïve IgG, TB IgG and TB αLy6G mice (n=5 mice per group, 3 independent experiments), paired one-way ANOVA. **l)** Quantification of organoid formation efficiency from intestinal epithelial cells isolated from naïve IgG, TB IgG and TB αLy6G mice (n=3 mice per group, 3 independent experiments), paired one-way ANOVA.

Considering the high epithelial turnover of the intestine, where the highly proliferative Lgr5⁺ stem cells can rapidly reshape the epithelial landscape, we performed PHATE analysis on the entire epithelial compartment of the intestinal Scrase datasets (Figure 2f, 2g and Extended Data Figure 2d). This allowed visualization of the diverse epithelial lineages, enterocytes, goblet, paneth, tuft and enteroendocrine cells, all of which originate from Lgr5+ stem cells (Figure 2g). Interestingly, we observed changes in the proportions of epithelial cell lineages in tumour-bearing mice, including an expansion of cluster 1 and a reduction in clusters 2 and 4 enterocytes, along with an increase in secretory lineages such as paneth cells (Extended Data Figure 2e and 2f). These changes were largely dependent on the presence of neutrophils (Extended Data Figure 2e and 2f). Moreover, these alterations extended beyond quantitative changes. MELD analysis demonstrated increased MELD likelihoods in cluster 1 enterocytes, goblet, and paneth cells within tumour-bearing, neutrophil-proficient mice relative to naïve or tumour-bearing neutrophil-depleted controls, reflecting significant transcriptional perturbations (Figure 2h-j and Extended Data Figure 2g). Accordingly, we observed an increased secretory lineage commitment in the transcriptional signature of Lgr5+ stem cells from tumour bearing mice in the presence of neutrophils (Extended Data Figure 2h and 2i).

Collectively, this analysis of epithelial cells in the lung and intestine revealed neutrophil-driven perturbations of progenitor cell states and lineage commitment in the presence of a distal primary tumour. This phenomenon is functionally linked to the enhanced organoid-forming efficiency previously observed (Figure 1b), as it is diminished in both the lung and intestine of tumour-bearing mice upon neutrophil depletion (Figure 2k and 2l).

### UPP1 is a present in neutrophils produced via emergency granulopoiesis

In the tumour-bearing host, neutrophils undergo a marked expansion in the bone marrow before entering the circulation and distant tissue where they mediate pro-metastatic and pro-tumorigenic activities^1,7,23,24^. This pro-tumorigenic, cancer-primed phenotype is observed by scRNA-seq in mature bone marrow neutrophils when neutrophil production shifts from steady-state to cancer-induced emergency granulopoiesis (Figure 3a). As expected, cancer-primed mature neutrophils exhibited an activated phenotype with elevated expression of multiple inflammatory pathways. Interestingly, this coincided with the activation of metabolic programs (Figure 3a). To further characterise the role of altered metabolism in acquisition of the cancer-primed state, we performed LC-MS/MS metabolomic analysis of media conditioned by splenic or lung neutrophils isolated from orthotopic mammary tumour-bearing, or naïve mice. Multiple tissue-depended changes were detected (Figure 3b). Focusing on specific cancer-dependent changes consistent across both tissues we focused on a switch from uridine to uracil release (Figure 3b). This was particularly evident when using the physiological, uridine-containing medium Plasmax, where only cancer-primed neutrophils consumed uridine and released uracil (Figure 3c and 3d). This pattern of uridine/uracil release points to the presence of Uridine Phosphorylase 1 (UPP1), which catalyses the phosphorolytic cleavage of uridine to uracil and ribose-1-phosphate (Figure3e). Indeed, cancer-primed neutrophils exhibit induction of UPP1 at the transcript level, accompanied by detectable protein level (Figure 3f and 3g). Most importantly, circulating neutrophils from treatment-naïve, newly diagnosed early breast-cancer patients^25^ displayed elevated UPP1 expression compared to their match healthy volunteers (Figure 3h). UPP1 enzymatic activity also reflected in an neutrophils-dependent uracil increase in the plasma of mice harbouring tumours as well as in their lung interstitial fluids (Figure 3i an Extended data figure 3a). Remarkably, similar trends in plasma uracil levels were detected in asymptomatic, newly diagnosed patients with early-stage breast cancer^25^ (Figure 3j).

**Figure 3.**
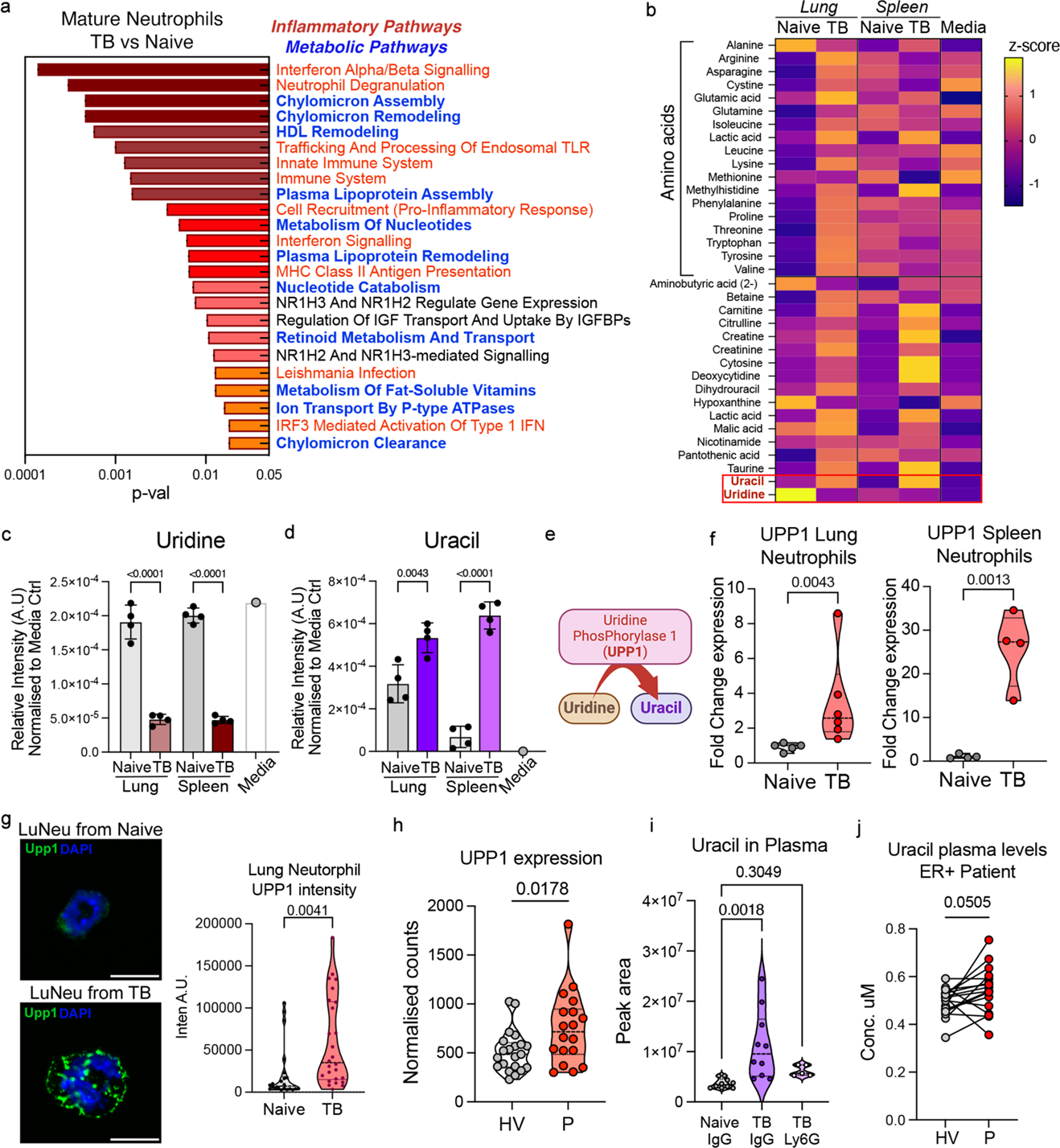
Cance-primed neutrophils acquire Upp1 expression and adopt an uracil secreting phenotype. **a)** Pathways enriched in mature bone marrow (BM) neutrophils from tumour bearing (TB) versus Naïve mice. Inflammatory and metabolic pathways are coloured in red and blue, respectively; enrichment P values are indicated. **b)** LC–MS–based metabolic profiling of control or neutrophil-conditioned media. Neutrophils were isolated from lung or spleen of naïve or TB (4T1 Balb/c) mice (n=4 per group) and incubated in Advanced DMEM/F12 medium. Data are shown as mean z-score per metabolite. **c-d)** Abundance of uridine (c) and uracil (d) in neutrophil-conditioned or control media generated from lung or splenic neutrophils isolated from naïve or TB (4T1 Balb/c) mice, incubated in uridine-containing Plasmax medium (n=4 mice per group), ordinary one-way ANOVA. **e)** Schematic representation of Uridine phosphorylase 1 (Upp1) enzymatic activity. **f)** Upp1 mRNA expression in lung and splenic neutrophils from naïve or TB (PyMT-FVB) mice lung (n=5 per group, spleen n=4 per group), two-tailed students t-test. **g) (Left)** representative image and **(Right)** quantification of UPP1 immunofluorescence intensity in lung neutrophils (LuNeu) isolated from naïve or tumour bearing (PyMT-FVB) mice. Data are represented a per cell mean intensity (n=4). Welch’s t-test. **h)** Plasma Uracil abundance from Naïve (Balb/c) IgG, TB (4T1) IgG or TB (4T1) αLy6G mice measured by LC-MS (Naïve IgG and TB IgG n=10, TB αLy6G n=5), ordinary one-way ANOVA. **i)** Upp1 expression in circulating neutrophils from healthy volunteers (HV) and patients with breast cancer (P). Data are normalized counts from bulk RNA-seq of purified neutrophils (n=14 per group), two-tailed Wilcoxon test. Hormone-receptor status of patients is indicated. **j)** Plasma uracil concentrations in matched samples from HV and P with breast cancer (n=17 per group), quantified by LC–MS, two-tailed Wilcoxon test.

This UPP1 expression in neutrophils in the context of breast cancer was recently reported as part of their pro-metastatic activity within the lung^26^. However, UPP1 expression also accompanies prolonged emergency granulopoiesis^27^, such as during bacterial infection^28^ or DSS colitis, but not more transient neutrophil responses following viral infection (Extended Data Figure 3b-e). Therefore, we next investigated how this fundamental UPP1 switch influences cancer-induced emergency granulopoiesis.

### UPP1 is a molecular switch required for the cancer-primed phenotype of neutrophils produced via emergency granulopoiesis

We used scRNAseq and PHATE analysis of neutrophils and their progenitors, from both Naïve and tumour-bearing mice, to dissect neutrophil differentiation trajectories from steady state to cancer-induced granulopoiesis (Figure 4a, 4b and Extended Data Figure 4a, 4b). Along the neutrophil differentiation trajectory, UPP1 upregulation was first detected at the immature neutrophil (Imm-Neu) stage and maintained in mature neutrophils (Figure 4c). To assess the potential role of UPP1 in this process, we analysed cancer-induced emergency granulopoiesis in UPP1-deficient mice^29^ and compared transcriptional perturbations across cellular states leading to neutrophil production using MELD analysis (Figure 4d). Strikingly, we observed marked UPP1-dependent perturbations not only in mature neutrophils but also at the level of the early progenitors (pro-neutrophils, Pro-Neu), which precede the proliferative progenitor pool (pre-neutrophils, Pre-Neu), (Figure 4d and Extended Data Figure 4c). This change was further supported by RNA velocity analysis, which showed that the boost in pro-neutrophil differentiation observed during the transition from steady-state (naïve WT) to cancer-induced emergency granulopoiesis (tumour-bearing WT) was absent in UPP1-deficient mice (Figure 4e, top panel and 4f). Nonetheless, the overall number of pro-neutrophils was maintained reflecting no changes in overall neutrophils produced between WT and UPP1-deficient mice harbouring tumour (Figure 4f and Extended Data Figure 4d and 4e).

**Figure 4.**
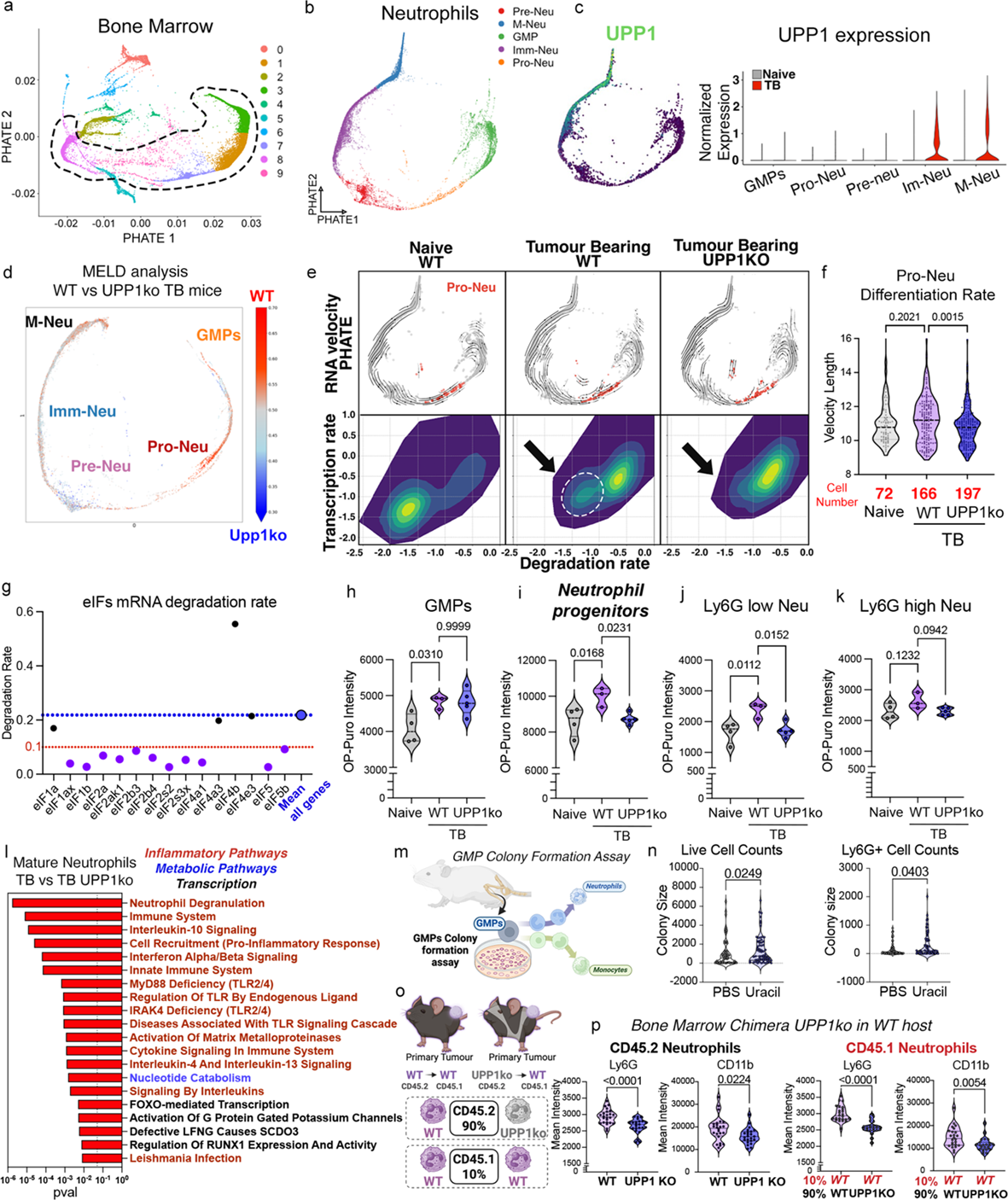
UPP1 is required to sustain the elevated translational rate of neutrophil progenitors and to enable the acquisition of the cancer-primed neutrophil phenotype. **a)** PHATE embedding of bone marrow (BM) scRNAseq data from Naïve WT (C57/Bl6), Tumour-bearing (TB) (PyMT-C57/Bl6) WT and TB UPP1KO (PyMT-C57/Bl6) mice. Cells are coloured by cluster. Neutrophil and neutrophil progenitor clusters are outlined with dashed line. **b)** Granulopoiesis neutrophil and progenitor clusters from (a) represented by PHATE embedding. Cells are coloured by neutrophil and progenitor subtype as indicated in the legend. **c)** UPP1 expression across the dataset, shown per cell on the PHATE embedding **(left)** and as normalized expression across neutrophil subclusters **(right)**. **d)** MELD transcriptional perturbation analysis showing per-cell relative likelihoods for TB WT versus TB UPP1KO samples. Red indicates higher likelihood associated with the TB WT condition. **e)** RNA velocity and transcriptional dynamics across pro-neutrophil (Pro-Neu) cells. **(Top)** RNA velocity vectors overlaid on PHATE embedding indicating differentiation trajectories within the granulopoiesis dataset; Pro-Neu population marked in red. **(Bottom)** Predicted transcription (Y-axis) and degradation (X-axis) rates for individual genes in Pro-Neu cells derived from scVelo dynamical modelling. Dashed lines and arrow encircle a group of ‘protected’ transcripts with low degradation rates specific to the TB WT condition. **f)** Velocity length as a measure of differentiation rate within Pro-Neu population from Naïve WT, TB WT and TB UPP1KO mice. Data represent average velocity length per cell within the Pro-Neu population. Cell numbers are indicated in red, Kruskall-wallace test. **g)** Predicted degradation rates of eukaryotic translation initiation factor (eIF) transcripts within the stable RNAs in Pro-Neu population from TB mice (highlighted in g). Mean degradation rate for all transcripts is indicated in blue and ‘protected’ transcripts with degradation rate < 0.1 are shown in purple. **h-k)** In vivo translation rate measured by OP-Puro incorporation in Naïve WT, TB WT and TB UPP1KO mice. OP-Puro fluorescence intensity was quantified by flow cytometry in GMPs (h), pre/proNeu (i) immature Ly6G low (j) and mature Ly6G high (k) bone marrow neutrophils isolated (Naïve WT n=4, TB WT n=3 TB UPP1KO n=5). Ordinary one-way ANOVA. **l)** Pathway enrichment analysis in mature neutrophils (M-Neu; from panel b) comparing TB WT and TB UPP1KO mice. Enrichment P values are shown. Inflammatory pathways are indicated in red, metabolic pathways in blue and transcription in black. **m)** Schematic of GMP colony-formation assay. FACS-sorted GMPs from naïve mice (FVB) were plated one cell per well with or without uracil. After colonies established, single GMP-derived colonies were profiled by FACS for cell number (colony size) and neutrophil/monocyte content (Ly6G+ and Ly6C+ cell count). **n)** Colony size **(left)** and Ly6G⁺ cell number per colony **(right)** from GMP colony-formation assay with or without uracil (n=6 mice. 62 PBS and 66 uracil-treated colonies) Mann-Whitney test. **o)** Schematic of bone-marrow chimera experiment. CD45.2 recipient mice (C57/Bl6) were reconstituted with either WT or UPP1KO CD45.1 bone marrow (C57/Bl6), generating chimeras with 10% WT CD45.1 and 90% WT or UPP1KO CD45.2 cells. This setup allows profiling of WT CD45.1 cells within a WT or UPP1KO haematopoietic environment. Mire where transplanted with PyMT tumour cells (C57/Bl6). **p)** Ly6G and CD11b intensity in circulating WT or UPP1KO CD45.2+ (donor) neutrophils and in WT CD45.1+ (recipient) neutrophils from bone marrow chimeric mice. (WT n=24, UPP1KO n=23 mice) Welch’s t-test.

Interestingly, detailed analysis of mRNA turnover during cancer-induced emergency granulopoiesis revealed that the pool of RNA transcripts normally stabilized within pro-neutrophils was specifically lost in UPP1-deficient mice. (Figure 4e, bottom panel and Extended Data Figure 4f and 4g). Among the pool of stable RNA transcripts in pro-neutrophils, we observed a notable enrichment of mRNAs encoding translation initiation factors (eIFs), suggesting a possible role for UPP1 in sustaining translational activity (Figure 4g and Extended Data Figure 4h-j). *In vivo* assessment of nascent polypeptide synthesis using O-propargyl puromycin (OP-Puro) labelling revealed a marked increase in translational activity from steady-state (naïve) to cancer-induced emergency granulopoiesis. Notably, this increase was absent in neutrophils from tumour-bearing UPP1-deficient mice (Figure 4h-k and Extended Data Figure 4k and 4l). The UPP1-dependent effect was absent in GMPs but was pronounced in neutrophil progenitors and remained evident along the neutrophil maturation trajectory (Figure 4h-k). Differential gene expression and pathway enrichment analyses of mature neutrophils from tumour-bearing WT and UPP1-KO mice indicated that these UPP1-driven alterations in mRNA stability, translation, and differentiation rates, although dispensable for increased neutrophil production, are essential for the full acquisition of the cancer-primed state (Figure 4l, see also Figure 3a for Naïve vs WT comparison).

In tumour-bearing mice, the UPP1-dependent alterations in Pro-Neu cells, occurred despite undetectable UPP1 expression (Figure 4c), suggesting a paracrine regulation by downstream UPP1-expressing neutrophil populations. We therefore tested whether uracil produced by immature and mature neutrophil could directly influence neutrophil production. Indeed, GMPs from naïve bone marrow, when allowed to differentiate ex vivo in the presence of uracil, showed a direct and specific increase in neutrophil output (Figure 4m, 4n and Extended Data Figure 4m). This suggests that a local increase in uracil, produced by UPP1-expressing immature and mature neutrophils, may modulate the behaviour of their upstream progenitors and establish a feedback loop that influences subsequent neutrophil production. To test this hypothesis, we generated WT or UPP1 KO bone marrow chimeric mice, where donor BM could be tracked using CD45.1/CD45.2 markers. The resulting chimeras showed ∼90% donor engraftment and 10% host cells >12 weeks post-transplant (Figure 4o). Chimeras were transplanted with breast cancer to activate cancer-induced emergency granulopoiesis. In line with the impaired acquisition of activated cancer-primed features, engrafted (CD45.2⁺) WT cancer-primed neutrophils displayed higher surface expression of Ly6G and CD11b compared with engrafted UPP1-deficient neutrophils (Figure 4p, left panel). Importantly, host derived (CD45.1⁺) WT neutrophils in UPP1KO chimeras, where uracil production is largely absent, similarly showed reduced Ly6G and CD11b expression, underscoring uracil’s role in shaping neutrophil phenotype (Figure 4p, right panel).

Collectively, these data establish UPP1-dependent uracil production as a feedback mechanism acting on neutrophil progenitors to determine neutrophil priming in the bone marrow. In the absence of UPP1, cancer-induced emergency granulopoiesis still drives robust neutrophil production; however, these cells fail to fully acquire the cancer-primed phenotype.

### UPP1 is required and pro-regenerative and pro-tumorigenic activity of cancer-primed neutrophils

To assess whether cancer-primed, UPP1-expressing neutrophils mediate the enhanced resilience to injury seen in the colon of tumour-bearing mice, we depleted neutrophils for five days following DSS administration. This intervention did not affect primary tumour growth but abrogated the tumour-associated reduction in colitis severity relative to naïve controls (Figure 5a and 5b and Extended data Figure 5a). To directly probe the role of UPP1 specifically, DSS colitis was induced in UPP1ko bone marrow chimeras (Figure 4o). No difference in primary tumour was observed (Extended data Figure 5b). While naïve animals showed no genotype-dependent difference in colitis response, tumour harbouring mice reconstituted with UPP1KO marrow exhibited greater injury severity than those with WT bone marrow (Figure 5c and 5d).

**Figure 5.**
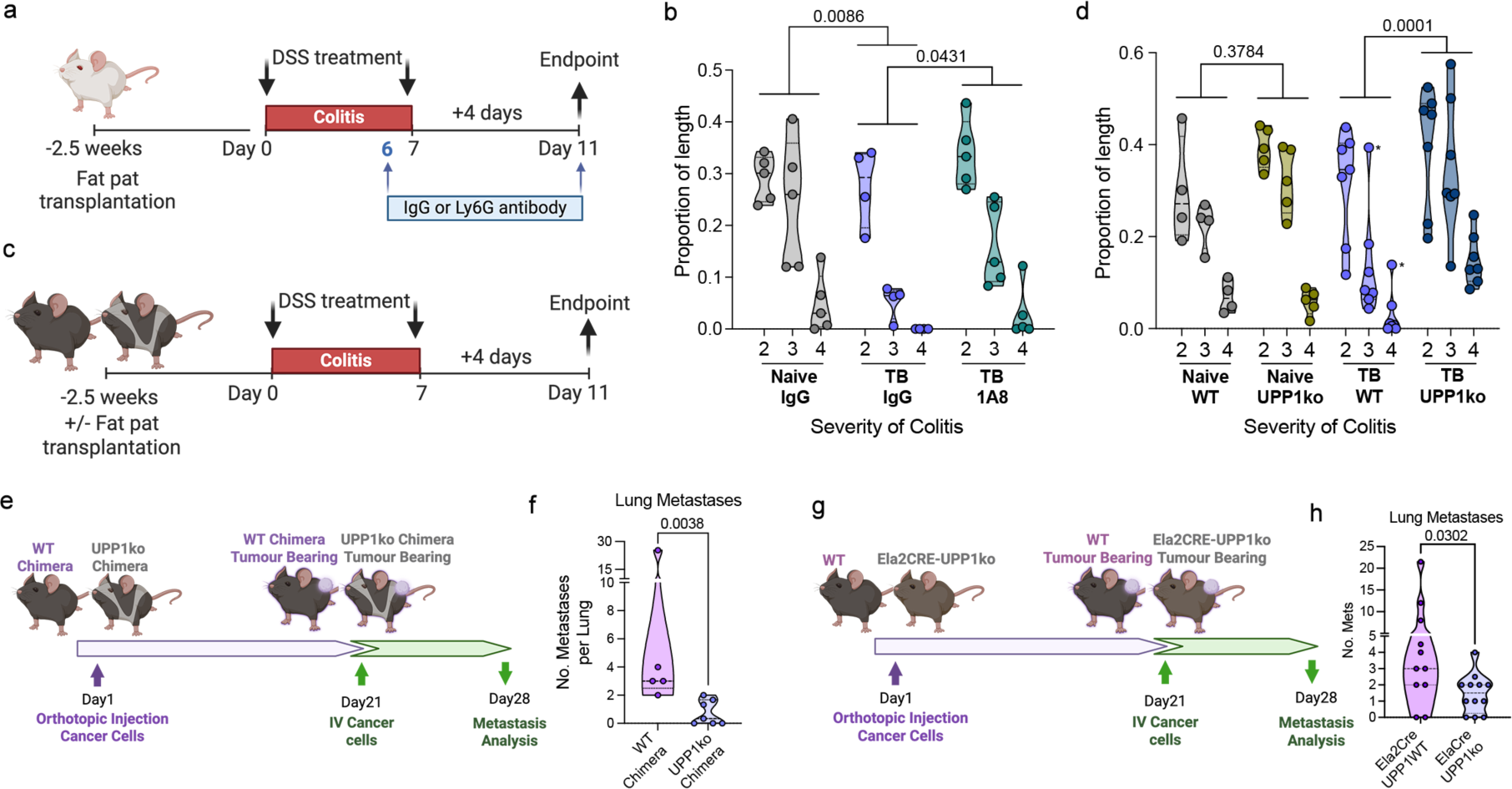
Tumour-bearing mice with UPP1-deficient neutrophils show reduced resilience to DSS-induced colitis and lung metastasis. **a)** Schematic for DSS induced colitis assessment in tumour bearing mice following neutrophil depletion. Naïve or MMTV-PyMT (FVB) orthotopic mammary tumour bearing animals received DSS drinking water (or control) for 7 days. From day 6 mice receive αLy6G antibody or isotype control (IgG). Colitis assessment was performed on day 11. **b)** Quantification of DSS-induced colitis severity in naïve IgG, tumour-bearing (TB) IgG and TB αLy6G (1A8) mice based on histological grading of colon sections. Data represent the proportion of colon length exhibiting each severity grade (n=5 mice per group), ordinary two-way ANOVA. **c)** Schematic of DSS-induced colitis in WT and UPP1KO bone-marrow chimeras. WT or UPP1KO chimeras were either left untreated or given DSS treatment for 7 days. Colitis severity was assessed 4 days after treatment. **d)** Quantification of DSS-induced colitis severity in naïve WT, naïve UPP1KO, TB WT and TB UPP1KO bone-marrow chimeras based on histological grading of colon sections. Data represent the proportion of colon length exhibiting damage severity (naïve WT n=4, naïve UPP1KO n=5, TB WT n=7 and TB UPP1KO n=7 mice), ordinary two-way ANOVA. **e)** Schematic for metastasis assessment in tumour-bearing UPP1KO bone-marrow chimeras. WT or UPP1KO BM-chimeras bearing orthotopic MMTV-PyMT tumours were intravenously injected with primary MMTV-PyMT tumour cells 7 days before assessing metastatic efficiency. **f)** Quantification of lung metastasis based on histological analysis (WT n=5, UPP1KO n=7), Mann-Whitney test. **g)** Schematic of metastasis assessment in WT (Ela2het, C57/Bl6) and neutrophil-conditional UPP1fox/flox (Ela2Cre-UPP1KO, C57/Bl6) mice. Mice bearing orthotopic MMTV-PyMT tumours were intravenously injected with primary MMTV-PyMT tumour cells (C57/Bl6) 7 days before assessing metastatic efficiency. **h)** Quantification of lung metastasis based on histological sections from tumour-bearing WT and Ela2CRE-UPP1KO mice. Data represent number of metastatic foci per lung (WT n=11, Ela2CRE-UPP1KO n=12), Mann-Whitney test.

The pro-metastatic phenotype of neutrophils can be directly demonstrated by the increased cancer cell lung colonization observed in naïve mice pre-conditioned with donor cancer-primed neutrophils (Extended data Figure 5c and 5d). When UPP1-KO cancer-primed neutrophils were used to pre-condition naïve lungs, they showed markedly reduced pro-metastatic conditioning activity compared to WT neutrophils (Extended data 5e and 5f). In line with a general decrease in pro-metastatic behaviour of cancer-primed UPP1KO neutrophils, a reduction in lung metastasis was observed in chimeras with UPP1KO compared to WT bone marrow (Figure 5e and 5f). UPP1 expression within the bone marrow is confined to neutrophils during emergency granulopoiesis (Extended Data Figure 3d), nonetheless, to test the specific deficiency of UPP1 in neutrophils, we generated conditional Upp1^flox^ mice and cross them with Ela2Cre transgenics to achieve neutrophil-specific UPP1 deletion. Consistent with the results from UPP1KO bone marrow chimeras, tumour-bearing Ela2Cre-Upp1^KO^ mice developed fewer lung metastases than Ela2Cre-WT controls, despite no change in primary tumour growth (Figure 5g, 5h and Extended Data Figure 5g).

These data collectively show that the deficient cancer-primed phenotype observed in UPP1-deficient mice (Figure 4l) results in a reduction of their pro-metastatic properties.

### Cancer-primed neutrophils promote platelet–AT2 cell proximity, which activate AT2 progenitor function and enhance tumour initiation

We next asked if the altered cancer-primed phenotype of tumour-bearing Ela2Cre-Upp1^KO^ mice influenced the activation of lung epithelial progenitors we have initially described (Figure 1 and 2). Indeed, alveolar cells from tumour-bearing Ela2Cre-Upp1^KO^ mice no longer showed enhanced organoid forming capacity compared to Ela2Cre-WT Naïve mice (Figure 6a and 6b).

**Figure 6.**
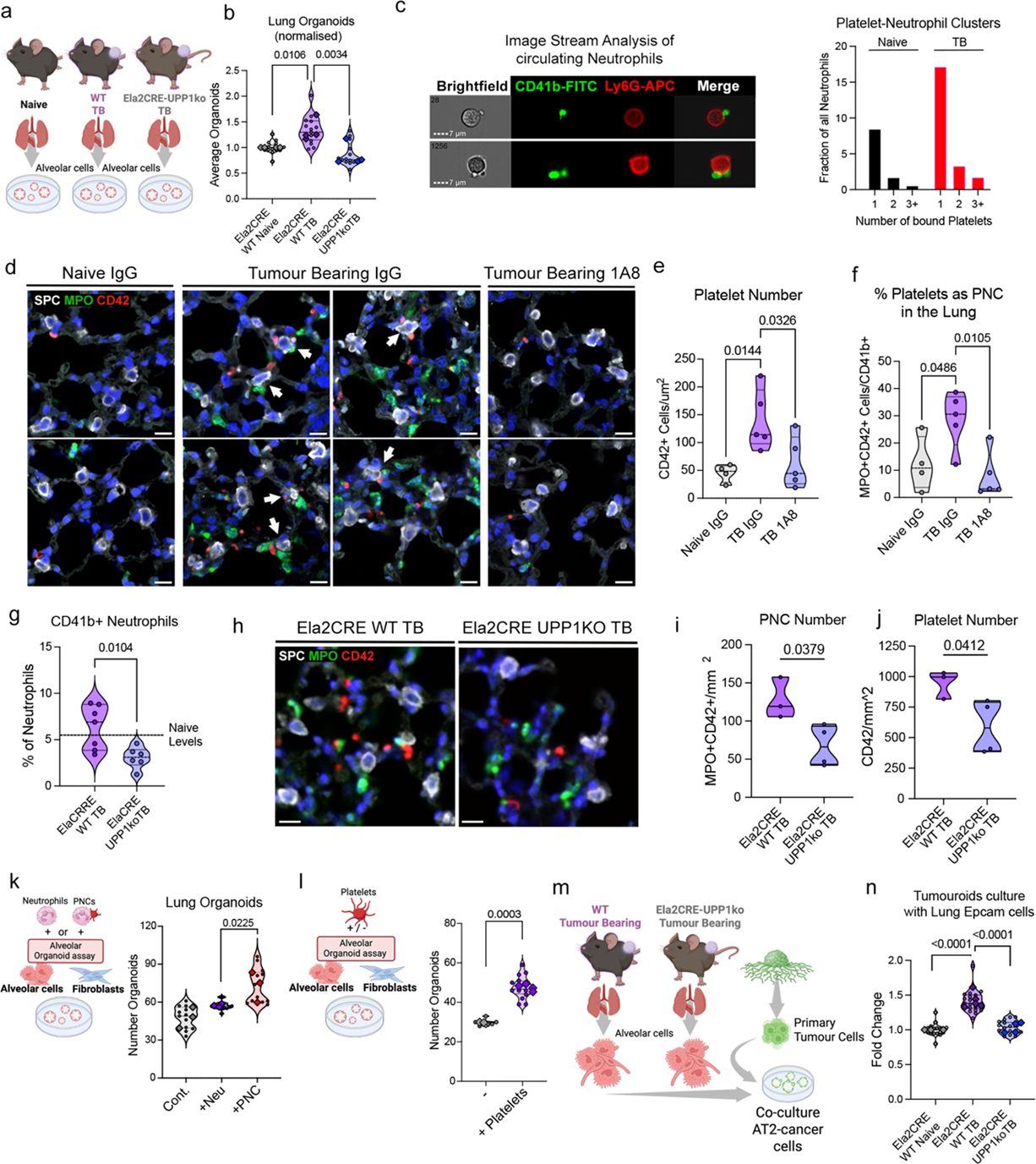
Cancer-primed neutrophils mediate a platelet dependent activation of AT2 cells. **a)**chematic of experimental design to assess lung epithelial organoid formation in mice with neutrophil specific UPP1KO TB mice. **b)** Quantification of organoid formation efficiency of lung epithelial cells isolated from naïve WT, tumour-bearing (TB) WT and TB Ela2CRE-UPP1KO mice (PyMT, C57/Bl6) (n=4 per biological replicates per group, large points, technical replicates, small points). one-way ANOVA performed on mean value of biological replicates. **c) (Left)** Representative brightfield and immunofluorescence images from ImageStream analysis of circulating neutrophils from tumour-bearing mice. CD41 (FITC, green) marks platelets and Ly6G (APC, red) marks neutrophils. **(Right)** Quantification of platelet-neutrophil clusters (PNCs) in circulation of a naïve (black) and a tumour-bearing (red) animals, shown as the percentage of PNCs containing increasing platelet numbers. **d)** Representative IF images of perfused lung from naïve IgG (FVB), TB (PyMT, FVB) IgG an TB (PyMT, FVB) αLy6G mice stained with SPC (AT2 cells, white), MPO (neutrophils, green) and CD42 (platelets, red). **e)** Quantification of platelet abundance in perfused lung from naïve IgG, TB IgG an TB αLy6G mice from (d), represented as platelets per mm2 of lung tissue. (naïve IgG n=4, TB IgG and TB αLy6G n=5), ordinary one-way ANOVA. **f)** Percentage of platelet-neutrophil clusters (CD42+ platelets in contact with MPO+ neutrophils) out of total platelets in lungs of naïve IgG, TB IgG an TB αLy6G mice (naïve IgG n=4, TB IgG and TB αLy6G n=5), ordinary one-way ANOVA. **g)** Percentage of circulating CD41b+ neutrophils out of all neutrophils in tumour-bearing Ela2CRE WT and Ela2CRE-UPP1KO mice (Ela2CRE WT n=7, Ela2CRE-UPP1KO n=6). (PyMT, C57/Bl6). Unpaired t-test. **h)** Representative IF images of perfused lung from TB Ela2CRE-WT and TB Ela2CRE-UPP1KO mice (PyMT, C57/Bl6), stained for SPC (AT2 cells, white), MPO (neutrophils, green) and CD42 (platelets, red). **i)** Quantification of platelet-neutrophil clusters represented as number of MPO+ neutrophils contacting CD42+ platelets per mm2 lung tissue. (Ela2CRE WT n=3, Ela2CRE-UPP1KO n=4) unpaired t-test. **j)** Quantification of platelet number (CD42+) per mm2 lung tissue in TB Ela2CRE-WT and Ela2CRE-UPP1KO mice from (h) (Ela2CRE WT n=3, Ela2CRE-UPP1KO n=4) unpaired t-test. **k) (left)** Schematic of experimental design and **(right)** lung epithelial organoid formation efficiency in presence of *ex vivo* generated platelet-neutrophil clusters (FVB) (+PNC), or neutrophils alone (+Neu) (n=3 biological replicates per group, large points, technical replicates, small points). Ordinary one-way ANOVA on mean of biological replicates. **l) (left)** Schematic of experimental design and **(right)** lung epithelial organoid formation efficiency in presence of platelets (+Platelets) (FVB) (-n=2, +Platelets n=4, biological replicates, large points, technical replicates, small points) Welch’s t test on mean of biological replicates. **m-n)** Schematic of experimental design (m) and tumour organoid formation efficiency (n) of primary MMTV-PyMT mammary tumour cells (C57/Bl6) cultured with or without lung epithelial cells isolated form naïve Ela2CRE-WT, TB Ela2CRE-WT or TB Ela2CRE-UPP1KO mice (C57/Bl6) (naïve Ela2CRE-WT n=4, TB Ela2CRE-WT n=5 or TB Ela2CRE-UPP1KO n=4, technical replicates shown in small points) Ordinary one-way ANOVA on mean of each n.

To identify microenvironmental signals downstream of neutrophils that drive alveolar progenitor activation in the pre-metastatic lung, we performed NicheNet ligand-receptor analysis^30^ on the expanded alveolar epithelial progenitor population defined by PHATE analysis of the entire lung epithelial compartment (Extended data figure 6a-d). This approach revealed a distinct pattern of neutrophil-dependent transcriptional perturbation within alveolar progenitors (Extended data figure 6d) and identified platelets as the cell type expressing the highest ranked ligand, Tgfβ1, predicted to drive this response (Extended Data Figure 6e). Because platelets and neutrophils are known to form stable interactions in circulation, and their presence is associated with poor prognosis in cancer patients^31^, we assessed whether platelet-neutrophil interactions were altered in the tumour bearing host. Because imaging flow cytometry confirmed platelet-neutrophil clusters (PNCs) formation in circulation of both naïve and tumour bearing host (Figure 6c), we next investigate their distribution within the lung. In perfused lung of naïve mice, under homeostasis, platelets reside within the lung microvasculature and do not directly contact alveolar cells (Figure 6d and 6e). However, we found that, in addition to neutrophils, platelets and PNCs were present within the lung interstitial space of perfused lungs from tumour-bearing mice, in close proximity to AT2 cells (Figure 6d-f and Extended Data Figure 6f and 6g). Importantly, depletion of neutrophils abrogated this effect, and neutrophil-dependent platelet presence in the lung interstitial space was supported by a direct correlation between neutrophil and platelet occurrence across different lung regions (Figure 6d-f and Extended Data Figure 6f-i).

In line with a deficient cancer-primed phenotype of UPP1KO neutrophils, which is also reflected in reduced expression of adhesion molecules (Figure 4p), we observed a decreased in circulating PNCs in Ela2Cre-Upp1^KO^ tumour harbouring mice (Figure 6g). Importantly, also in perfused lungs of Ela2Cre-Upp1^KO^ tumour harbouring mice, both platelets and PNCs were reduced (Figure 6h-j and Extended Data Figure 6j), indicating a reduction of this extravascular platelet displacement. Importantly, co-culture of lung epithelial cells with *ex vivo*-generated PNCs showed that only platelet-decorated neutrophil, but not neutrophils alone, enhanced alveolar organoid formation (Figure 6k). This indicates that epithelial cells which would normally lack progenitor activity to initiate organoids acquire this potential in the presence of PNCs. Notably, platelets alone were also able to enhance epithelial organoid formation (Figure 6l).

We have previously shown that co-culture with AT2 cells directly enhances organoid formation by primary PyMT breast cancer cells^13^. We therefore asked whether these activated AT2 cells might provide a potential advantage to cancer cells within pre-metastatic lungs. Remarkably, AT2 cells isolated from tumour-bearing mice significantly enhanced PyMT tumour organoid formation in direct correlation with their regenerative capacity, whereas AT2 cells from Ela2Cre-Upp1^KO^ tumour harbouring mice, which lacked progenitor activation, did not promote tumour organoid initiation (Figure 6m, 6n). A similar correlation was observed when UPP1 deletion was driven by a second neutrophil-specific promoter (S100A8Cre) (Extended Data Figure 6l, 6m). Collectively, these data suggest that cancer-primed neutrophils facilitate platelet infiltration into the lung parenchyma of tumour-bearing hosts, and that platelet-derived signals contribute to the activation of a pro-regenerative program in lung alveolar cells, further supporting the establishment of a pre-metastatic niche.

## Discussion

Primary tumours condition distant organs long before metastatic cells arrive, releasing systemic factors and altering immune cell dynamics to create permissive pre-metastatic niches^1,2^. Traditionally viewed as organ-specific and deterministic, cancer-induced conditioning, when mediated by tumour-driven systemic inflammation, can instead be seen as part of a broader, and perhaps more fundamental, physiological process, one that cancer co-opts to systemically alter the homeostatic state of distant organs. Many features of the metastatic niche mirror those that accompany regenerative responses following injury^9^. Given that pre-metastatic conditioning initiates perturbations in distant organs that are subsequently exacerbated within the local metastatic niche^1^, it is plausible that the underlying cancer-induced systemic program co-opts regenerative pathways. Indeed, our previous work showed that activation of epithelial regenerative programs characterise the early metastatic niche and enhance cancer initiation potentia^l13,16^. We therefore asked whether an epithelial regenerative response is a component of this cancer systemic conditioning.

Using organoid initiation efficiency as a measure of epithelial progenitor activation, we found enhanced organoid formation in tumour-bearing mice, not only in organs typically targeted for metastasis such as lung and liver, but also in pancreas and intestine, which are rarely metastatic targets. This systemic increase in regenerative activity altered how tissues respond to acute injury. When lung damage was induced by influenza infection in tumour-bearing mice, the extent of post-infection injury was markedly reduced compared naïve controls, despite similar severity of infection. A similar phenomenon was observed in the colon following DSS-induced colitis. Whether this reduced tissue damage in mice harbouring distal tumours reflects protection from the initial insult or an expedited repair process, it demonstrates that tumour-induced changes reprogram steady-state physiology towards increased resilience to an injury event.

We and others have shown that neutrophils responses to sterile injury can support tissue regeneration^16,32,33^. Consistently, single cell transcriptomic analysis and functional organoid formation assessment, revealed that neutrophils are important drivers of epithelial progenitor perturbation in the lung and intestine in tumour-bearing mice. In the cancer context, neutrophils are mobilised in high numbers through reprogrammed granulopoiesis, generating pro-tumorigenic or “cancer-primed” neutorphils^1,23,24^. This enhanced production represents a shift from steady state to emergency granulopoiesis^27^. To characterise the cancer-primed state of neutrophils we performed single cell transcriptomic profiling of bone marrow neutrophils in naïve and tumour-bearing mice. Given that many metabolic pathways were altered, we assess neutrophil metabolic activity by performed an unbiased metabolomic analysis of media conditioned by control and cancer-primed neutrophils isolated from lungs and spleens. Focusing on changes occurring in neutrophils from both organs, we found that cancer-primed neutrophils generated a strong accumulation of uracil, at the expenses of uridine. This metabolic switch reflected the activity of Uridine Phosphorylase 1 (UPP1) that catalyses the conversion of uridine to uracil. UPP1 expression was detected exclusively in immature and mature neutrophils when production occurred via emergency granulopoiesis, whether tumour induced or triggered by other stimuli.

Analysis of neutrophils from UPP1KO tumour-bearing mice revealed a profound alteration of their cancer-primed phenotype, without alterations in granulopoiesis output. Within bone marrow, we detected a disruption in the differentiation trajectory in the pro-neutrophil pool, a stage where UPP1 is not expressed, suggesting a non-cell autonomous effect. This impaired trajectory correlated with increased turnover of transcripts enriched for translation initiation factors, indicative of a potential reduction in translational activity. Indeed, functional labelling of nascent proteins showed that, while granulocyte-monocyte progenitors (GMPs) show a cancer-induced increase in translation independent of UPP1, neutrophil progenitors and their immature neutrophil progeny showed an UPP1-dependent increase. We found that exogenous uracil influenced *ex vivo* production of neutrophils from isolated GMPs and *in vivo*, reduction of uracil levels in UPP1KO bone marrow chimeras, attenuated acquisition of cancer primed phenotype in the small pool of host derived WT neutrophil. We therefore postulate that uracil produced locally by immature neutrophils establishes feedback loop targeting neutrophil progenitors, to enable the full acquisition of the cancer-primed state during elevated granulopoiesis. These data not only define a new mechanism required for the phenotypic integrity of cancer-primed neutrophils, but, as UPP1 expression is induced during infection-driven emergency granulopoiesis, could also be important to study the neutrophil responses more broadly.

We found that enhanced resilience to damage induced by DSS-colitis in tumour-bearing mice was dependent on UPP1, despite no effect in naïve animals. Moreover, the altered cancer-primed phenotype of UPP1KO neutrophils blunted their pro-metastatic activity. Part of this deficient phenotype may also reflect the absence of UPP1 itself as its expression in lung neutrophils was recently shown to support metastasis via uracil-induced extracellular matrix remodelling^26^. Here, we observed that the activation of epithelial regeneration represents and additional mechanism of pre-metastatic conditioning whereby epithelial cells in an active regenerative state directly support tumour cell initiation *ex vivo*. Mechanistically, we found that cancer-primed neutrophils activate alveolar progenitors via their interactions with platelets. Neutrophils-platelet clusters were shown to be important mediators of inflammatory responses^34^ and their characteristic transcriptional signature was shown to be enriched in circulation of cancer patients with the poor prognosis^31^. While in the lung, platelets normally contribute to alveolar repair following injury^35^. In the cancer context, they have been implicate in promoting metastasis^36–38^, including recent evidence for a key role of platelet activity in high fat diet mediated pre-metastatic lung conditioning^39^. Here, we show that tumour-bearing mice display a neutrophil-dependent re-localisation of platelets within the lung interstitial space, positioning them proximal to AT2 alveolar cells. By doing so neutrophils recapitulate platelet-induced alveolar progenitor response typically displayed upon an injury. Indeed, we could observe a direct platelet-induced alveolar progenitor activation in *ex vivo* co-culture, and neutrophils gained this capacity when decorated with platelets. Importantly, we found a significant reduction in platelet accumulation within the lung interstitial space of tumour-bearing UPP1KO mice, accompanied by reduced epithelial progenitor activity and loss of the ability of these cells to boost tumoroid initiation.

Collectively, our findings show that cancer-induced systemic conditioning involves the activation of regenerative programs in distant tissues, altering their physiological state. In organs targeted for metastatic growth, this enhanced epithelial progenitor function can directly boost tumour-initiating potential. We demonstrate that, in the lung, this phenomenon is mediated by cancer-primed neutrophils, which engage platelets to promote their interaction with alveolar cells. Furthermore, we identify UPP1 expression as an important regulator of neutrophil production during tumour-induced emergency granulopoiesis. In this context, UPP1 expression in immature/mature neutrophils supports uracil synthesis, optimal mRNA dynamics and the increased rate of translation needed for maintaining proper differentiation trajectories under conditions of accelerated neutrophil production. Elevated uracil was previously found in plasma of advanced breast cancer patients^26^, yet here we show that can be also detected in asymptomatic breast cancer patients at diagnosis^25^. Circulating neutrophils from newly diagnosed cancer patients also show an increased UPP1 expression, suggesting that this metabolic switch is active early in the tumorigenic program. This work suggests that targeting UPP1, could represent a potential strategy to disengage pro-metastatic neutrophils in cancer patients.

## Materials and Methods

### Mouse Strains

All mice were bred and maintained under specific-pathogen-free conditions by The Francis Crick Biological Research Facility and wildtype female mice were used between 6 and 14 weeks of age. Wild-type BALB/cJ, C57BL/6J, FVB/nJ mice from The Jackson Laboratory were provided by The Francis Crick Biological Research Facility. MMTV–PyMT mice are on either a FVB/NJ or C57BL/6 background. *Ela2-Cre* knock-in mice were originally purchased from the European Mouse Mutant Archive. *Mrp8-Cre* mice (B6.Cg-Tg(S100A8-cre,-EGFP)1Ilw/J, stock:021614) and *CD45.1* (B6.SJL-Ptprc^a^Pepc^b^/BoyJ, stock:002014) mice were purchased from Jackson and maintained on a C57BL/6J background. *Upp1KO* mice were purchased from the Mutant Mouse Resource and Research Centre (stock:037119-UCD) and maintained on a C57BL/6J background.

*Upp1-flox* mice were generated for us by Ozgene and maintained on a C57BL/6J background. Neutrophil specific Upp1 deficient mice were generated by crossing *Ela2-Cre* knock-in or transgenic *Mrp8-Cre* with *Upp1-flox* animals. Breeding and all animal procedures were performed at the Francis Crick Institute in accordance with UK Home Office regulations under project licenses P83B37B3C and PP5920580.

### Bone-marrow Chimeras generation

To generate BM chimeras, *CD45.1* C57BL/6J recipient mice were lethally irradiated (2 x 6 Gy, 15-hour interval) and reconstituted with 5 x 10^6^ bone marrow cells from WT or *Upp1 KO* C57BL/6J donor mice. BM chimeric mice were then left for 9 weeks before tumour cell injection to reconstitute hematopoietic compartment.

### Orthotopic Tumour cell injections

MMTV-PyMT cells were isolated from MMTV-PyMT tumours as previously described^40^. For orthotopic transplantation 0.75-1×10^6 cells, from either FVB/NJ or C57BL/6J background, were injected in 50ul growth factor reduced, phenol red free Matrigel (Corning) into one or both of the abdominal mammary glands (4^th^) under brief isoflurane anaesthesia in WT FVB/NJ, WT C57BL/6, *Upp1 KO, Ela2-Cre*, *Ela2Cre-Upp1flox*, BM Chimeric *CD45.1*-*WT/Upp1KO, Mrp8-Cre or Mrp8-Cre-Upp1flox*. With the exception of BM chimeric mice, animals were injected at 6-14 weeks of age.

### Metastasis Assay

For metastatic seeding assays, 1 x 10^6^ C57BL/6J MMTV-PyMT or FVB/NJ MMTV-PyMT, or 400k 4T1-GFP cells were injected in PBS into the tail vein of *Ela2-Cre*, *Ela2Cre-Upp1flox*, BM Chimeric *CD45.1*-*WT/Upp1KO or* BALB/cJ, animals either bearing orthotopic fat pad tumour or naive. With the exception of BM chimeric mice, animals were injected at 6-14 weeks of age. Lungs were harvested approximately 6 weeks after orthotopic mammary fat pad injection and fixed overnight in 10% NBF, prior to processing and embedding in paraffin. Histological quantification of tumours was done from H&E stained sections.

### *In vivo* Neutrophil Depletion

For neutrophil depletion in FVB/NJ and BALB/cJ mice, rat anti-Ly6G antibody (BioXcell, clone 1A8, 25ug per mouse) in PBS or rat IgG control (Cell Sciences Unit, Francis Crick Institute) was administered daily by IP injection.

### Cell Culture

All cell lines were provided by the Cell Services Unit of The Francis Crick Institute, where they were authenticated using Short-Tandom Repeat profiling and species-identification tests and confirmed to be mycoplasma-free. Cells were cultured in Dulbecco’s modified eagle medium (DMEM) (Fisher Scientific) supplemented with 10% foetal bovine serum (FBS; Fisher Scientific) and 100 U ml^−1^ penicillin-streptomycin (Fisher Scientific). MMTV-PyMT cells were isolated from late-stage carcinomas and cultured on collagen-coated dishes in MEM medium (DMEM/F12 (Fisher Scientific) with 2% FBS, 100 U ml^−1^ penicillin-streptomycin, 20 ng ml^−1^ EGF (Fisher Scientific) and 10 μg ml^−1^ insulin (Merck Sigma-Aldrich)). Collagen solution contains 30 μg ml^−1^ PureCol collagen (Advanced Biomatrix), 0.1% bovine serum albumin and 20 mM HEPES in HBSS (Fisher Scientific). All cells were cultured at 37 °C and 5% CO_2_.

### Tissue Digestion for cell isolation or analysis

Lung tissue was minced with scissors and digested with Liberase TM and TH (Roche Diagnostics) and DNase I (Merck Sigma-Aldrich) in HBSS -Ca-Mg (1.5 ml digestion mix per lung) for 30 min at 37 °C in a shaker at 180 r.p.m. Samples were passed through a 100 μm filter and centrifuged at 1,250 r.p.m. for 10 min.

Bone marrow (BM) was isolated from the leg bones (tibia/femur). The bones were first flushed with HBSS -Ca,-Mg (Fisher Scientific), 10% FCS before crushing. The cell preparation was filtered through 100 μm filter and centrifuged at 1,250 r.p.m. for 10 min. Spleen were bisected and crushed through a 100 um cell strainer and rinsed through with HBSS -Ca,-Mg (Fisher Scientific), 10% FCS, before centrifugation at 300 x g. for 10 min.

Lung, bone marrow and spleen cell-pellets were incubated in Red Blood Cell Lysis buffer (Miltenyi Biotec) for 5 min at room temperature and quenched with 10% FCS HBSS -Ca-Mg and passed through a 70 um strainer. After centrifugation, cells were washed with magnetic-activated cell sorting (MACS) buffer (0.5% BSA and 250 mM EDTA in PBS) and passed through a 20 μm strainer-capped tube to generate a single-cell suspension. Antibody staining was then performed for cell isolation (flow cytometry or magnetic-bead based cell separation) or for flow cytometry analysis.

Hepatocytes were prepared from mice by two-step collagenase perfusion. Briefly, after placing catheter into the portal vein, the inferior vena cava was cut and the liver was perfused at 5–7 ml/min with pre-warmed Liver Perfusion Medium (Fisher Scientific) for 10 minutes. Then the liver was perfused with pre-warmed Digestion Medium, DMEM low glucose (1g/ml), Type IV collagenase 0.6g/l at 5 ml/min for 3–5 minutes. After dissociation, cells were filtered through a 70 μm filter and enzyme activity quenched with ice cold DMEM+10% FCS. Hepatocytes were further separated and purified by centrifugation at low speed (50 g, 1–3min) and Percoll gradient centrifugation, to remove dead cells and debris.

For acinar cell preparation, pancreas was carefully dissected and collected in ice cold HBSS -Ca,-Mg (Fisher Scientific). Pancreas were minced with scissors and digested in 20mL of digestion medium composed of Advanced DMEM/F12 (Fisher Scientific) and 1mg/mL collagenase type V from Clostridium histolyticum (Sigma-Merck). Digestion was performed for 20 minutes in a 37℃ shaker at 140rpm. The reaction was stopped by adding 30mL of ice-cold MACS buffer containing (0.5% BSA and 250 mM EDTA in PBS). To obtain a single cell suspension, samples were centrifuged for 4 min at 400 x g at 4℃, and the pellet was resuspended in MACS buffer and filtered twice with a 70 um and 40μm filters sequentially.

### Fluorescence-activated cell sorting analysis (FACs)

Single-cell suspensions were incubated with mouse FcR Blocking Reagent (Miltenyi Biotec) for 10 min at 4 °C followed by incubation with fluorescently conjugated antibodies (prepared in PBS with fixable Zombie live/dead stain) for 20 min at 4 °C. Cells were washed twice with MACS buffer, prior to fixation. Countbright^TM^ absolute counting beads (Thermo Scientific) were used for absolute cell sample counts. Flow cytometry analyses were performed on a BD LSR-Fortessa (BD Biosciences), BD X20 (BD Biosciences) or BD Symphony (BD Biosciences) and subsequently analysed using FlowJo v.10.4.2 (FlowJO, LCC).

### Magnetic cell sorting (MACS)

EpCAM^+^ lung epithelial cells or Ly6G^+^ neutrophils were isolated from mouse single-cell suspensions via MACS. Cells were incubated with FcR Blocking Reagent for 10 min at 4 °C.. For EpCAM^+^ isolation, cell preparations were first incubated with anti-CD45-beads and anti-CD31beads (Miltenyi Biotec) for 15 min at 4 °C, washed and then negatively sorted through LS columns (Miltenyi Biotec) to deplete CD45+ and CD31+ cells from the sample. Flow through were then incubated with anti-EpCAM beads for positive selection through LS/MS magnetic separation columns. For neutrophil isolation, cell preparations were first incubated with anti-Ly6G FITC conjugated antibody (1:100) for 15 min on live, washed and then incubate wth with anti-FITC beads (Miltenyi Biotec). After washing with MACS buffer, positive cell selection was performed using LS/MS magnetic separation columns (Miltenyi Biotec). Neutrophil pellets sorted for RNA isolation were resuspended in Qiazol Lysis Reagent (Qiagen), snap frozen on dry ice and stored until isolation.

### Organoid Formation assays

Organoids were prepared from single cell preparations (as described above) from naïve or tumour bearing FVB/NJ or C57BL/6J mice.

#### Lung Organoid

Lung epithelial cells (EpCAM^+^CD45^−^CD31^−^) were isolated as described above by MACS and were resuspended in lung organoid media (DMEM/F12 supplemented with 10% FBS, 100 U mL–1 penicillin-streptomycin, 1 mM HEPES, GlutaMAX™ and insulin-transferrin-selenium (ITS) (Merck Sigma-Aldrich). For each technical replicate, 7k Epithelial Cells were mixed with 35k murine normal lung fibroblast (MLg) cells and resuspended in 100 μl organoid media and GFR Matrigel (1:1 ratio) before plating in 24-well polystyrene transwell insert, 0.4 μm pore size (Corning). After incubating for 30 min at 37 °C, 500 μl organoid media was added to the lower chamber and media changed every two days. Bright-field images were acquired after 14 days using an EVOS microscope (Thermo Fisher Scientific) and quantified using FiJi (ImageJ).

#### Intestinal Organoid

Intestinal organoids were generated from freshly isolated mouse intestine adapted from Gracz e.t al 2012^41^. Proximal intestine was opened longitudinally, and villi were scraped off with a coverslip. Intestines were then incubated in PBS containing 30 mM EDTA, 1.5 mM DTT and 10uM Y27632 StemCell Technologies for 20 min on ice before being transferred to PBS containing 30 mM EDTA and 10uM Y27632 at 37 °C for 10 min. To release epithelial cells, the tissue was shaken vigorously before removal of remaining intestinal tissue. Crypts were washed and filtered through a 70 um strainer. Crypts were resuspended in Advanced DMEM F/12 containing 10mM HEPES, 100 U/ml penicillin-streptomycin and 1:100 Glutamax, counted to normalise crypt density per sample and plated in Cultrex BME Type 2 RGF Pathclear (3533-01002; Amsbio). Organoids were maintained in an in-house generated media composed of Advanced DMEM F/12, 50ng/ml EGF, Noggin-CM and RSPO1-CM, 1% B27, 1.25mM NAC, 10mM Nicotinamide and 10 um Y27632. 5-7 days after plating, once crypt cultures had established, crypts were collected in cold Advanced DMEM F/12, mechanically dissociated using glass pipette, washed and counted and seeded in Culltrex BME at 200 crypts per droplet and maintained in ENR media as described above without Nicotinamide or Y27632 addition. Organoid number was assessed 7 days post seeding using an EVOS microscope (Thermo Fisher Scientific). RSPO-1 and Noggin conditioned media were generated by Cell Sciences, Francis Crick Institute).

#### Liver Organoid

1 x 10^4^ hepatocytes were seeded in a 25 μl drop of GFR Matrigel in 24 well low attachment plates. Organoids were maintained with media changes every 2-3 days of Advanced DMEM containing: pencillin/streptomycin (1:100), Glutamax (1:100), 10 mM HEPES, B27-Vitamin A (1:50), 10 mM Nictotinamide, 10 nM Gastrin I, 50 ng/ml EGF (Peprotech), 50 ng/ml FGF10 (Peprotech), 25 ng/ml HGF (Peprotech), 50ng/ml FGF7 (Peprotech), 1.25 mM N-acetylcysteine, RSPO1-CM (15%-Cell Sciences, Francis Crick Institute), 3 μM CHIR99021 (Sigma), 1 μM A83-01 (2BScientifc) and 10 uM Y27632 (StemCell Technologies) as reported in (https://doi.org/10.1016/j.cell.2018.11.013). The number of organoids in each well were quantified 14-18 days post seeding using an EVOS microscope (Thermo Fisher Scientific).

#### Pancreatic Organoid

Freshly dissociated acinar cells were counted manually and 1×10^4^ acinar cells were seeded in 25μl drop of growth factor reduced Matrigel in 24 well low attachment plates. Organoids were maintained Advanced DMEM containing: pencillin/streptomycin (1:100), 50 ng/ml EGF (Peprotech), 100 ng/ml FGF10 (Peprotech), 10 nM Gastrin I (Sigma), 10% Noggin-CM (Cell Sciences, Francis Crick Institute), 10% RSPO-CM (Cell Sciences, Francis Crick Institute), 1.25 mM N-acetylcysteine (Sigma), 10 mM Nicotimanide (Sigma), 1X B27 Supplement (Fisher Scientific), as adapted from (DOI 10.1038/emboj.2013.204). Media were changed every 2-3 days. The number of organoids in each well were quantified 10 days post seeding using an EVOS microscope (Thermo Fisher Scientific).

### Tumour organoid co-culture assay

To assess tumour organoid formation in co-culture with lung epithelial cells, GFP+ MMTV-PyMT cancer cells were thawed and seeded on collagen-coated plates. The following day, mouse lung EpCAM+ cells were isolated from either naive or orthotopic MMTV-PyMT mammary tumour-bearing FVB/NJ mice, or from naive or orthotopic MMTV-PyMT mammary tumour-bearing WT Ela2Cre^+/−^ or Upp^fl/fl^ Ela2Cre^+/−^ animals by MACS as described above. GFP+ MMTV-PyMT cancer cells were detached using PBS/EDTA and trypsin and were filtered through a 20 μm cell strainer to obtain a single-cell suspension. For each technical replicate, 1 × 10^4 lung EpCAM+ cells were mixed with 1 × 10^3 GFP+ MMTV-PyMT cancer cells, centrifuged, and resuspended in a 1:1 mixture of GFR Matrigel with Lung Organoid Media to a final volume of 100 μl. This mixture was plated in transwell inserts, and 500 μl of Lung Organoid Media was added to the base of each well. 7 or 14 days following plating, tumour organoid number was assessed using brightfield and fluorescence imaging with an EVOS microscope.

### Lung organoid formation with PRP and ex vivo generated PNC

To isolate platelet-rich plasma (PRP), fresh blood was collected by cardiac puncture from FVB/NJ mice into citrate buffer–containing tubes. Blood was diluted to a final volume of 3 mL in Wash Buffer (modified Tyrode’s buffer - 134 mM NaCl, 3 mM KCl, 0.3 mM NaH_2_PO_4_, 5 mM HEPES, 5 mM Glucose, 2 mM MgCl_2_, 1.2 mM NaHCO_3_, 0.2% w/v BSA, 1 mM EGTA pH 6.8) and centrifuged at 200 x g for 5 min at room temperature with low acceleration and no brake. To isolate platelets, the PRP-containing supernatant was collected and centrifuged at 800 x g for 10 min with low brake. The resulting platelet pellet was resuspended in Resuspension Buffer (identical to Wash Buffer but lacking EGTA, adjusted to pH 7.2).

Primary lung EpCAM⁺ cells were isolated from FVB/NJ mice by MACS sorting, as described above. For each technical replicate, EpCAM⁺ lung epithelial cells and MLg fibroblasts were centrifuged and resuspended in a 1:1 mixture of growth factor–reduced (GFR) Matrigel and Lung Organoid Medium, supplemented with or without platelets at a final concentration equivalent to 20% of that found in the circulation. Lung organoid formation was assessed after 7 days, as described above

### Platelet-neutrophil cluster generation and organoid co-culture

Platelets were isolated as described above and activated by resuspension in DMEM/F12 media supplemented with 10 nM freshly thawed phorbol 12-myristate 13-acetate (PMA), at a concentration equivalent to that found in the circulation. Platelets were incubated at 37 °C for 30 min and washed twice at 800 × g (low acceleration, low brake) to remove residual PMA. Lung neutrophils were isolated from 6–8-week-old FVB/NJ mice by MACS, as described above, and incubated with activated platelets at 37 °C for 1 h to generate platelet–neutrophil clusters (PNCs). Neutrophils and PNCs were washed twice at 300 × g to remove unbound platelets, Purified PNCs or untreated neutrophils were then co-cultured with lung EpCAM⁺ epithelial cells and MLg fibroblasts for the lung organoid formation assay, as described above. For each technical replicate, a 20:1 PNC-to-EpCAM⁺ cell ratio was used. Lung organoid formation was quantified after 7 days.

### Influenza induced lung injury

Influenza virus strain X31 (H3N2-supplied by John Cauley, WHO Collaborating Centre for Reference and Research on Influenza, Francis Crick Institute) were generated in allantoic cavity of 10-day-embryonated hen’s eggs and were free of bacterial, mycoplasma, and endotoxin contamination. All viruses were stored at −80°C and titrated on Madin–Darby canine kidney (Mdck) cells as plate forming units (PFU).

Naïve C57BL/6J or MMTV-PyMT tumour bearing animals were infected intranasally with 50ul of PBS containing 250 PFU of X31virus under light isoflurane anaesthesia. Mice were monitored daily for weight loss. Mice were 8-14 weeks old at time of infection and were monitored daily for body weight and clinical symptoms. Pre-infection body weights were recorded, and tumour-bearing mice were weighed regularly to correct for tumour mass when calculating infection-induced weight loss.

### DSS induced colitis

Acute colitis was induced in FVB/NJ, BM Chimeric *WT/Upp1KO* mice, either naïve or bearing orthotopic MMTV-PyMT tumours, by administering 2% Dextran Sulphate Sodium (DSS) salt, colitis grade (Fisher Scientific, 12871781) in drinking water for 7 days, followed by plain water. Pre-colitis body weights were recorded, and mice were weighed daily throughout until weight recovered post-treatment. Clinical symptoms, including diarrhoea and rectal bleeding, were monitored throughout. Tumour-bearing mice were weighed regularly, and tumour volume measurements were used to correct body weight changes associated with colitis. With the exception of BM chimeric mice, DSS induction was performed in animals aged 8–14 weeks. At the experimental endpoint, intestines were harvested, and faecal material was removed by gentle flushing with 50 ml ice-cold PBS using a syringe fitted with a gavage needle. Tissues were fixed flat between blotting paper in 10% neutral-buffered formalin for histological assessment.

### Tissue Fixation and Histology

Mouse tissues were fixed overnight in 10% neutral-buffered formalin (NBF) and transferred to 70% ethanol processing and embedding in paraffin blocks. Serial sections (4-5 μm) were cut using a microtome (Leica) and mounted onto Superfrost Plus slides. Section levels were spaced 100-150 μm apart. Slides were deparaffinized and rehydrated using standard methods.

### Immunohistochemistry

FFPE sections were baked for 1 h at 60°C before staining. Immunohistochemistry was performed using a Roche Ventana Discovery automated stainer. For Ki67 detection (Abcam, Ab15580 1:6000), antigen retrieval was carried out for 48 min at 90°C with CC1 solution followed by detection with DISCOVERY Omnimap anti-Rabbit HRP (Roche, 05269679001). For S100A9 (in house 2B10 clone, 1:2500), antigen retrieval was carried out for 8 mins with Roche Protease I at 37°C and detection used DISCOVERY Omnimap anti-Rat HRP (Roche 05891892001). Slides were counterstained with haematoxylin and coverslipped using a Tissue-Tek Prisma Plus automated stainer. Stained slides were scanned on a Zeiss AxioScan.Z1 digital slide scanner and analysed quantitatively using QuPath software.

### Immunofluorescence

#### Upp1 staining of Neutrophils

MACs sorted Ly6G+ neutrophils (as described above) were mounted onto poly-lysine coated coverslips (VWR) at a density of 1 x 10^6^/ml in PBS at 37 °C for 30 minutes. Slides were then fixed with 4% PFA for 15 minutes at room temperature prior to permeabilization with PBS 0.2% Triton X-100, washing with IF Wash Buffer (PBS, 7.5mM NaN_3_, 0.1% BSA, 0.04% Tween-20, pH 7.4) and blocking with IF Wash Buffer + 3% BSA. Primary antibody Upp1 (Proteintech 14186-1-AP, 1:200) in IF Wash Buffer 1% BSA was incubated overnight at 4 °C prior to washing with PBS. Secondary antibody (Alexa Fluor 488 donkey anti-rabbit IgG, 1:250) and DAPI was prepared in PBS/1% BSA and incubated at room temperature for 2 hours prior to washing and mounting. Images were obtained with Zeiss Invert 880 with Airyscan and quantified using QuPath.

#### Spc, Mpo, CD41b Triple-Immunofluorescence of Lung

Lungs were obtained from naïve and MMTV-PyMT tumour bearing animals perfused with PBS to remove blood from the tissue. Lungs were then dissected, fixed, processed and embedded as described above.

Triple Immunofluorescence staining was performed on the Leica Bond Rx automated stainer. MPO (R&D, AF3667 1:400), CD42b (Abcam, ab183345 1:300), and SFTPC (Abcam, ab211326 1:5000) were sequentially stained and detected with Immpress anti-Goat-HRP (Vector, MP-7405-500) or Novolink Max Polymer (Leica, RE7260-CE) and Opal TSA reagents (Akoya FP1487001KT Opal 520, FP1488001KT Opal 570, FP1497001KT Opal 690). Antigen retrieval with ER2 at 95°C for 20minutes was performed before each primary antibody to remove the antibody complex. Slides were counterstained with DAPI (Thermo Scientific, 62248) 1:2500 and mounted with Prolong Gold Antifade reagent (Invitrogen, P36934).

Images were obtained with Zeiss Invert 880 with Airyscan and quantified on QuPath.

### Image flow cytometry

Blood was collected by cardiac puncture into EDTA-containing tubes and 100 ul was added to 1 ml of pre-cooled PBS containing 1% PFA. At least 1 hour following fixation, samples were quenched in 5 ml FACS buffer before proceeding with standard RBC lysis and FACS antibody staining with anti-Ly6G-APC (1A8, Biolegend) and anti-CD41b-FITC (MWReg30, Biolegend) as described above.

Samples were acquired with a Amnis ImageStream MkII imaging flow cytometer (Cytek) equipped with 405nm, 488nm, 561nm and 642nm excitation lasers. CD41b -FITC emission was collected in Channel 2: 480-560nm, Ly6G-APC emission was collected in channel 11: 660-740nm. Images were collected using a 60x objective lens. Data were analysed using IDEAS software, version 6.4.8 (Cytek)

### In vivo Neutrophil conditioning

Ly6G+ neutrophils were MACS sorted from lungs of orthotopic 4T1 tumour-bearing BALB/c mice, or from orthotopic MMTV-PyMT tumour-bearing C57BL/6J WT or whole-body Upp1 KO animals generated as described above. Following lung tissue harvest and enzymatic digestion (as described above), single cell suspensions were incubated with FITC-conjugated anti-Ly6G (1A8, BioLegend) antibody before positive selection by MACS with anti-FITC-conjugated magnetic beads (Miltenyi Biotec). For experiments involving C57BL/6J WT and Upp1 KO mice, lungs from 2 mice were pooled per sample. Isolated neutrophils were resuspended in filtered PBS, and 1.5 × 10^6^ cells (from 4T1 BALB/c donors) or 8 × 10^5^ cells (from pooled MMTV-PyMT C57BL/6J donors) were filtered through 35 µm cell-strainer FACS tubes (Falcon, blue cap) immediately before injected intravenously into each naive recipient mouse. Transfers were performed three times over the course of one week. One day after the final transfer, recipient mice were intravenously injected with either 0.4 × 10^6^ 4T1-GFP+ or MMTV-PyMT BL6 primary mammary tumour cells, filtered immediately prior to injection. For experiments involving 4T1-GFP+ cancer cells, metastatic burden was assessed by FACS analysis of GFP+ cells in the lung as a proportion of CD45-negative cell abundance. In the case of MMTV-PyMT BL6 metastases, lungs were collected 2 weeks following intravenous injection, fixed in 4% PFA, and embedded in paraffin. Metastatic burden was calculated from 4 μm serial sections stained for haematoxylin and eosin (H&E) by manually counting the total number of metastatic foci using a Zeiss Axio Scan.Z1.

### qPCR

Cell populations were homogenised using Qiazol lysis reagent followed by phase separation, as described in the Qiazol Handbook (!REF) (Qiagen). Upper aqueous phase was transferred to a fresh tube before addition of 70% Ethanol. RNA was then isolated using RNeasy Micro kit according to manufacturer’s instructions. cDNA was synthesized using random primers, using Superscript III Frist Strand Synthesis kit (Thermo) according to manufacturer’s recommendations. qPCR was performed using Powerup SYBR Green Master Mic (Thermo) and QuantStudio 3 RT-PCR System (Thermo). hypoxanthine phosphoribosyl transferase 1 (*Hprt1*) was used.

### O-propargyl-puromycin (OP-Puro) Assay

Cellular translation rate was measured *ex vivo* in OP-Puro administered mice ex vivo as described in^42^. Briefly, WT C57BL/6j and UppKO C57BL/6j fat pad injection of 1×10^6 MMTV-PyMT BL6 cells to generate orthotopic fat lad tumours. 10mM OP-Puro in PBS was administered at 10ul/g of mouse weight. An additional cohort of naïve mice were also administered with PBS (10ul/g) for OP-Puro negative controls. OP-Puro administration was staggered, and mice were culled and bone marrow was harvested exactly 1 h after OP-Puro/PBS administration. Bone marrow was processed for FACS staining as described above and stained with Lineage Cocktail, anti-CD16/32, anti-CD34, anti-CD117, anti-CD115, anti-CD11b, anti-Ly6C, anti-Ly6G, anti-CD184. Detection of incorporated OP-Puro was performed using Click-iT reaction with Alexa Fluor 555 (Thermo Fischer), according to manufacturer’s guidelines.

### GMP colony formation and phenotyping assay

Methocult 3434 methocellulose medium containing 100 u/ml penicillin-streptomycin with and without 1 mM Uracil was prepared and added to flat bottomed 96-well dishes. Bone marrow cells were isolated from tibia and femur of 6-8 week old FVB NJ animals (two mice pooled per sample) as described above. Following RBC lysis, and washing, cells were stained with GMP panel comprising lineage markers (CD3, CD4, CD8, B220, Ly6C, Ly6G), Ly-6A/E, CD117, CD16/32, CD34.

GMPs (live lin-, c-Kit high, Sca1 neg, CD34+ CD16/32+) were sorted using a BD Influx Cell sorter (BD Biosciences) with plate sorting function and plated one cell per well in methocellulose-containing plates. GMP-derived colonies were left to grow for 11 to 14 days before individual colonies were harvested and stained for Lineage cocktail (anti-CD3, anti-CD4, anti-CD8, anti-B220, anti-CD19, anti-NKp46), anti-Ly6A/E, anti-CD117, anti-CD16/32, anti-CD34, anti-CD184, anti-Ly6C, anti-CD11b, anti-Ly6G, and fixed. Individual colony size, monocyte and neutrophil populations were assessed by FACS using a A ZE5 Cell Analyzer (Bio-Rad). Colonies with high LSK positive cell proportion were omitted from analysis.

### Single-cell RNA sequencing (scRNAseq) Lung

Female FVB/NJ mice (6–8 weeks old) were injected orthotopically with 1×10^6 primary MMTV-PyMT tumour cells to induce mammary tumours. Three weeks after tumour induction, tumour-bearing animals and age-matched naïve controls received daily intraperitoneal injections of either IgG or anti-Ly6G antibody (clone 1A8) for one week prior to tissue harvest. Lungs were collected and digested as described above, with the digestion time extended to 40 minutes to ensure efficient recovery of stromal cell types. Single-cell suspensions from two mice per condition were pooled and stained for CD45 and EpCAM to enable prior to FACS sorting. Cells were sorted sequentially such that 1/3 of the sample comprised each of CD45⁺ (immune), EpCAM⁺ (epithelial), and CD45⁻EpCAM⁻ (stromal/other) fractions. The number of each population was adjusted such that immune, epithelial, and stromal fractions each comprised approximately one-third of the total sample for downstream single-cell library preparation.

For each sample, an aliquot of cells was stained with the AO/PI Cell Viability Kit (Logos Biosystems) and counted using a LUNA-FX automated cell counter (Logos Biosystems). Cell viability prior to loading exceeded 80% for all samples. Single-cell suspensions were processed following the manufacturer’s instructions using the Chromium Next GEM Single Cell 3′ Reagent Kit v3.1 (10x Genomics). Approximately 10,000 cells per sample were loaded into Chromium Single Cell 3′ v3 chips for partitioning and barcoding. Libraries were prepared according to the manufacturer’s protocol and sequenced on an Illumina NovaSeq 6000 platform using paired-end reads.

Raw sequencing data were processed with Cell Ranger (v6.0.1, 10x Genomics) using the mm10-2020-A Mus musculus reference transcriptome (10x Genomics). The pipeline demultiplexed reads by cell barcode, aligned sequences using the integrated STAR aligner (v2.7.x), and generated filtered feature-barcode matrices excluding GEMs containing ambient RNA from lysed or low-quality cells.

Gene-barcode matrices from naïve IgG, PT IgG, and PT aLy6G libraries were imported into Seurat as individual objects. Features detected in fewer than three cells and cells with fewer than 200 detected genes were excluded. Data were processed using the standard Seurat workflow. Per-cell metrics, including mitochondrial and ribosomal transcript fractions, were calculated. During quality control, cells were retained if they contained 600–5,500 detected genes and <10% mitochondrial RNA. Each sample was log-normalised, and the 2,000 most variable features were identified using the vst method. After scaling, dimensionality reduction was performed using the first 30 principal components (PCs), followed by UMAP embedding (PCs 1–30) and Louvain clustering (resolution range 0.1–2.0). A resolution of 0.5 was selected for single-sample visualisation.

Broad cell-type identities were assigned based on canonical marker expression. Clusters with mixed lineage signatures were flagged as potential doublets and further assessed using scater and scDblFinder. Doublet classifications were recorded in the Seurat metadata, and only singlets were retained for downstream analysis.

For three-sample integration, singlet objects were SCTransformed with regression of percent mitochondrial gene expression. A total of 3,000 integration features were selected, and datasets were integrated using SCT normalisation. PCA was then performed using 30 PCs, followed by UMAP embedding (PCs 1–30) and clustering across resolutions (0.1–2.0). A resolution of 0.6 was used for visualisation.

Initial lineage annotation was guided by RNA-level expression of canonical markers including Epcam (epithelial), Sftpc (AT2), Ager (AT1), Krt8/18, Ptprc (immune), and Pecam1 (endothelial). The Epcam⁺Sftpc⁺ subset containing alveolar type II epithelial cells was saved as a separate Seurat object for downstream analyses.

For PHATE analysis, expression matrices and corresponding metadata were exported from Seurat. For pairwise MELD comparisons, Sftpc⁺ cells were further subset into Naïve IgG vs PT IgG and PT IgG vs PT aLy6G groups prior to export. Low-dimensional embeddings were generated using PHATE (as implemented by Moon, K. R. et al. 2019)^18^. Briefly, AT2 subset matrices were imported into Python and processed using the scprep package. Rare genes, as well as mitochondrial and ribosomal transcripts, were filtered out. Data were library-size normalised and square-root transformed, followed by PCA reduction. Two-dimensional PHATE embeddings were then computed using knn = 10 and decay = 15, with all other parameters left as default.

MELD (Burkhardt et al. 2021)^22^ was applied to estimate, for each cell, the relative likelihood of belonging to the perturbed versus control condition. The built-in parameter search was used to optimise k (nearest neighbours) and β (smoothing) by simulating perturbations on the same graph and selecting the (k, β) pair that minimised the mean-squared error between the ground-truth conditional density and the MELD estimate. MELD relative likelihoods were calculated separately for each pairwise comparison (Naïve IgG vs PT IgG; PT IgG vs PT aLy6G) using the optimised parameters.

To identify transcriptional perturbation within the expanded population of epithelial progenitors, expression matrices for epithelial cells were exported for PHATE analysis to capture the epithelial lineage hierarchy, using the same parameters (knn = 10, decay = 15). MELD was then applied as above to estimate per-cell likelihood scores corresponding to neutrophil depletion (αLy6G) or tumour-bearing (IgG) conditions. To define class categories for downstream differential gene expression analysis, MELD likelihoods were modelled using a three-component Gaussian mixture model with scikit-learn. Cells were classified according to their most probable Gaussian component, representing low, no, or high perturbation likelihood. Differential gene expression analysis was performed between class categories using diffxpy and EnrichR was used for pathway enrichment analysis, identifying transcriptional programs associated with epithelial progenitor activation in a neutrophil-dependent manner.

NicheNet (Browaeys,et a., 2020)^30^ was used to identify microenvironmental components predicted to explain the neutrophil-dependent transcriptional perturbation in alveolar progenitors (Receiver) observed tumour-bearing mice (PT IgG vs PT αLy6G conditions). All cell types from tumour-bearing IgG dataset were included as ligand-sources (Sender) to identify putative ligand–receptor interactions and upstream signalling pathways contributing to the observed alveolar progenitor transcriptional program. A default background gene detection (>10% per cluster) was used for Sender populations. For the receiver population, the top predicted ligands were ranked by Pearson correlation between ligand–target regulatory potential scores and target gene expression and ranked ligand expression levels were assessed across cell type clusters.

### Bone Marrow

WT or Upp1⁻/⁻ C57BL/6J mice received two mammary fat pad injections of 1 x 10^6^ primary MMTV-PyMT tumour cells to establish orthotopic mammary tumours. Four weeks after transplantation, bone marrow (BM) cells were harvested and processed for flow cytometry as described above, then stained using the BM enrichment panel for CD45, Ly6G, Ly6C, cKit, CD31. To ensure adequate representation of neutrophils and haematopoietic progenitors, populations were sorted sequentially to obtain approximately one-third myeloid cells (live CD45+, Ly6G+, Ly6C+), one-third haematopoietic progenitor cells (live CD45⁺cKit⁺), and one-third total bone marrow. Sorting was performed into BSA-coated collection tubes using a BD Influx cell sorter (BD Biosciences).

Single-cell suspensions from bone marrow were processed and sequenced using the same 10x Genomics Chromium Single Cell 3′ v3 workflow described above for lung datasets.

Gene–barcode matrices were imported into Seurat (v4.3) to generate individual Seurat objects. Data processing followed the standard Seurat workflow, as described for the lung dataset, with the same quality control thresholds except for an extended upper limit of 7,000 detected features per cell. Cells were retained if they contained >600 features and <10% mitochondrial RNA. Each sample was log-normalised, and variable feature selection, scaling, PCA (30 PCs), UMAP embedding, and Louvain clustering (resolutions 0.1–2.0) were performed as described above for the lung dataset.

Samples were then integrated using Seurat’s standard log-normalisation workflow. Briefly, filtered and normalised objects were scaled, integration features were selected, anchors were computed, and an integrated object was generated. The integrated data were subsequently processed for PCA reduction, UMAP embedding, and Louvain clustering as above.

Cell-type identities in the integrated bone marrow dataset were assigned using SingleR (Bioconductor), with reference transcriptomes from celldex. Canonical markers (Ptprc, Csf1r, Csf3r, and Ly6g) were visualised on the UMAP to guide annotation.

The integrated Seurat object was converted to a SingleCellExperiment object using DietSeurat. SingleR was run using multiple reference datasets (Monaco Immune, MouseRNAseq, and ImmGen), and the ImmGen reference was selected for final annotation. Main and fine cell-type labels were computed, with low-confidence assignments excluded. Cluster-level annotations were refined by providing Seurat cluster IDs to SingleR and manually curating the assignments based on marker expression.

Low-dimensional embeddings were generated using PHATE as described above for the lung dataset. PHATE embeddings were calculated on the full BM dataset with parameters knn = 20 and decay = 15 (remaining parameters set to default). Unsupervised clustering was performed using PHATE’s built-in clustering function (knn = 10). Cluster-wide expression of neutrophil and neutrophil progenitor markers was used to identify relevant clusters, which were subset for targeted downstream analysis. PHATE embeddings were recomputed for this neutrophil/progenitor subset before applying MELD to compute relative likelihoods for each pairwise comparison (WT Naïve vs WT PT and WT PT vs Upp1⁻/⁻ PT).

Aligned BAM files for each sample were processed using velocyto to generate a merged loom file containing spliced and unspliced gene counts. RNA velocities were computed using scVelo, run separately for each sample. Data were normalised and log-transformed (min_shared_counts = 20, n_top_genes = 500). Moments were calculated with parameters method = ‘umap’, n_pcs = 30, n_neighbors = 30. Gene-specific kinetic parameters were estimated using the dynamical model, and values for transcription (α), splicing (β), and degradation (γ) rates were extracted from the fit_alpha, fit_beta, and fit_gamma columns in the scVelo output matrices.

### Intestine

Female FVB/NJ mice (6–8 weeks old) were injected orthotopically with 1×106 primary MMTV-PyMT tumour cells twice to induce 2 bilateral mammary tumours. Three weeks after tumour induction, tumour-bearing animals and age-matched naïve controls received daily intraperitoneal injections of either IgG or anti-Ly6G antibody (clone 1A8) for 11 days prior to tissue harvest.

Intestines were dissected, flushed with ice-cold PBS, opened longitudinally, and the luminal surface was gently scraped with a coverslip to enrich for intestinal stem and progenitor compartments. Tissue was then incubated in PBS containing 30 mM EDTA, 1.5 mM DTT, and 10 μM Y-27632 (StemCell Technologies) for 20 min on ice, followed by incubation in PBS containing 30 mM EDTA and 10 μM Y-27632 at 37 °C for 10 min. Samples were shaken vigorously, and remaining intestinal tissue was removed. The released cells were washed and resuspended in Advanced DMEM/F12 supplemented with penicillin–streptomycin (1:100), GlutaMAX (1:100), 10 mM HEPES, and 1 mg/ml dispase/collagenase. Suspensions were incubated for 12 min at 37 °C with agitation every 2 min, then quenched with 10% FCS. DNase

I was added before sequential filtration through 70 μm and 40 μm strainers. Cells were pelleted and resuspended in PBS for single-cell library preparation.

Intestine single-cell suspensions were processed and sequenced using the same 10x Genomics Chromium Single Cell 3′ v3 workflow described above for lung datasets Gene–barcode matrices were imported into Seurat (v4.3) to generate individual objects for quality control and downstream analysis, following the same workflow used for the lung dataset. Cells were retained if they contained 700–7,000 detected features. Each sample was log-normalised, and the 2,000 most variable features were identified using the vst method. Data were scaled and reduced by PCA (30 PCs), followed by UMAP embedding (PCs 1–30) and Louvain clustering across resolutions 0.1–2.0. Visualisations were generated at the resolution used for the lung dataset.

Samples were integrated using Seurat’s log-normalisation anchor workflow (SelectIntegrationFeatures, FindIntegrationAnchors, IntegrateData), as described for the bone-marrow dataset. Broad lineages were assigned using canonical markers, after which epithelial clusters were subset for downstream analysis.

Raw expression and metadata matrices for epithelial subsets were exported, and low-dimensional embeddings were computed using PHATE as described for the lung dataset, with parameters knn = 13 and decay = 10 (other parameters set to default). MELD relative likelihoods were calculated as for the lung analysis, using the in-built parameter search function to optimise k and β values. MELD was run for each pairwise comparison, and resulting likelihoods were visualised on the PHATE embeddings.

Within epithelial subsets, differentially expressed genes (DEGs) were identified using Seurat’s FindMarkers() function. Genes were considered differentially expressed if expressed in >10% of cells in the positive group, with an average log2 fold-change >0.5 and adjusted p < 0.05. Pathway enrichment analysis was performed using Enrichr^43–45^.

### Published scRNAseq datasets

Single-cell RNA-seq datasets of mouse bone marrow and spleen^28^ were downloaded from the Gene Expression Omnibus (GEO) and processed using Seurat. Data processing followed the same standard workflow described above. Broad cell types were identified based on canonical marker gene expression. Upp1 expression patterns were assessed and visualised using the FeaturePlot() and VlnPlot() functions.

### Bulk RNA sequencing

Bulk RNA sequencing data from human neutrophil pellets (as described in Ramessur, A. et al. 2023)^25^ were sequenced on an Illumina NovaSeq 6000 platform targeting approximately 25 million reads per sample (50 bp paired-end read length). Bulk RNA-seq data were processed using the nf-core/rnaseq pipeline (version 3.10.1) implemented in Nextflow (version 23.10.0) with Singularity containers. Reads were aligned to the Homo sapiens reference genome (GRCh38, Ensembl release 95) using the RSEM–STAR option within the pipeline to generate gene-level count matrices.

Downstream analysis was performed in R using DESeq2 for differential expression analysis after filtering genes with fewer than ten total counts across samples.

### Software and Package versions

Unless mentioned explicitly in the text, all computational analyses were performed in R (v4.3.3) and Python (v3.9.18) and the following packages and libraries were used:

#### R environment

Seurat (v5.2.1), sctransform (v0.4.2), SeuratObject (v5.1.0), uwot (v0.2.3), scater (v1.30.1), scDblFinder (v1.16.0), SingleR (v2.4.1), celldex (v1.12.0), SingleCellExperiment (v1.24.0), Matrix (v1.6.5), nichenetr (v2.0.6), enrichR (v3.4), DESeq2(1.42.1).

#### Python environment

phate (v1.0.11), scprep (v1.2.3), meld (v1.0.2), scvelo (v0.3.1), velocyto (v0.17.17), loompy (v3.0.7), numpy (v1.26.3), pandas (v2.0.3), anndata (v0.10.3), scanpy (v1.9.6), umap-learn (v0.5.4), matplotlib (v3.8.0), seaborn (v0.12.2).

### Metabolomics analysis

#### Neutrophil conditioned media isolation for LC–MS

Lung and splenic neutrophils were isolated by MACS from either Naïve or 4T1 orthotopic fatpad tumour bearing Balbc/j mice as described above. Neutrophils were resuspended with serum-free 3D growth media containing DMEM/F12 supplemented with 100 U/ml penicillin-streptomycin, 20 ng/ml EGF, 20 ng/ml bFGF, 4 μg/ml Heparin and B27 (1:500) to a final cell density of 1×10^7 cells/ml. 100ul of Neutrophil suspension was into flat-bottomed ultra-low attachment 96-well plates along with neutrophil-free media controls. 16-18 h post plating, control and conditioned media were harvested on ice and centrifuged at 4°C at 300xg for 10 min to remove remaining cells and then at 100xg for 10 min to remove cell debris. Media samples were then frozen on dry ice and stored at -80°C prior to LC-MS/MS based metabolomics.

#### Lung interstitial fluid (LIF) isolation for LC–MS

Female BALB/c mice received single mammary fatpad injections of 5×10^5 4T1 cells to induce orthotopic mammary tumour growth. After 19 days, tumour-bearing mice, or naïve controls received daily intraperitoneal injections of anti-Ly6G (1A8) or isotype control antibody for 3 days prior to tissue harvest.

Mice received terminal anaesthetic under schedule 1 procedures and 400 ul saline containing 10 uM deuterated proline were delivered intratracheally as an internal standard for normalisation. Lungs were excised, blotted, weighed sliced in half and placed on pre-cooled Costar Spin-X Centrifuge Tube Filters (Corning). Samples underwent sequential centrifugation at 1,000, 2,000, and 4,000 rpm for 10 min each at 4 °C. The eluate was transferred to low-bind tubes and centrifuged at 16,000 × g, 5 min, 4 °C. Supernatants were aliquoted, snap-frozen, and stored at −80 °C.

For LC–MS extraction, 4 volumes of −20 °C 80% methanol were added to LIF, samples were vortexed, incubated 10 min at −20 °C, and centrifuged (16,000 ×g, 10 min, 4 °C). Supernatants were dried (SpeedVac) and reconstituted in instrument solvent. Signal normalisation recovered deuterated proline in each sample.

#### Neutrophil conditioned media and plasmax metabolite extraction

In summary, 5 µL of neutrophil conditioned media was transferred to a 1.5 mL Eppendorf tube, and mixed with 140 µL of water, 150 µL methanol and 50 µL chloroform, vortexed and centrifugated for 10 min at 4 °C 12700 rpm. A biphasic portioning was conducted, and the polar fraction was collected, dried, resuspended in 100 µL of methanol:water (1:1 v/v) containing 5 µmol L^-1^ of ^13^C_5_,^15^N_1_-valine and transferred to LCMS vials to analyse on the LCMS method. The plasmax metabolite extraction followed the same protocol described above.

#### Human plasma metabolite extraction for quantification of dihydrouracil, uridine, and uracil

A total of 20 µL of human plasma were mixed with 60 µL of MeOH, vortexed and centrifugated for 10 min at 4 °C 12700 rpm. Then, 55 µL of supernatant was collected and dried under N_2_ evaporator. Then, samples were resuspended in 50 µL of water, vortexed, transferred to LCMS vials and analysed on LC-MS/MS method. Qualitative and quantitative analysis was performed using XCalibur QualBrowser and Tracefinder 5.1 software.

#### Calibration curve extraction for quantification of dihydrouracil, uridine, and uracil

An internal calibration curve was prepared following the sample preparation described in the “metabolite extraction” section. A stock solution of 1.6 µmol L^-1^ of dihydrouracil, uracil and uridine were prepared in methanol, and 60 µL was added to 20 µL of human serum (Sigma Aldrich, H4522). In the end, a serial dilution was applied to obtain final concentrations of 1, 0.5, 0.2, 0.08, 0.04, 0.016, 0.008, 0.0032, and 0.0016 µmol L^-1^ of dihydrouracil, uracil and uridine. Three quality controls were prepared at the final concentration of 0.0048, 0.25, and 0.7 µmol L-1 using the same stock solution. Results were processed using GraphPad Prism 10.6.0 software.

#### LCMS method

Metabolite analysis was performed by LC-MS using a Q-EXACTIVE Plus (Orbitrap) mass spectrometer (ThermoFisher) coupled with a Vanquish UHPLC system (ThermoFisher). The chromatographic separation was performed on a SeQuant® Zic®pHILIC (Merck) column (5 μm particle size, polymeric, 150 x 4.6 mm). The injection volume was 5 μL, the oven temperature was maintained at 25°C, and the autosampler tray temperature was maintained at 4°C. Chromatographic separation was achieved using a gradient program at a constant flow rate of 300 μL/min over a total run time of 25 min. The elution gradient was programmed as decreasing percentage of B from 80 % to 5 % during 17 minutes, holding at 5 % of B during 3 minutes and finally re-equilibrating the column at 80 % of B during 4 minutes. Solvent A was 20 mM ammonium carbonate solution in water supplemented by 4 mL/L of a solution of ammonium hydroxide at 35% in water and solvent B was acetonitrile. MS was performed with positive/negative polarity switching using a Q-EXACTIVE Plus Orbitrap (ThermoFisher) with a HESI II probe. MS parameters were as follows: spray voltage 3.5 and 3.2 kV for positive and negative modes, respectively; probe temperature 320°C; sheath and auxiliary gases were 30 and 5 arbitrary units, respectively; and full scan range: 65–975 m/z with settings of AGC target and resolution as balanced and high (1e6 and 70,000), respectively. Data were recorded using Xcalibur 4.2.47 software (ThermoFisher). Mass calibration was performed for both ESI polarities before analysis using the standard ThermoFisher Calmix solution. To enhance calibration stability, lock-mass correction was also applied to each analytical run using ubiquitous low-mass contaminants. Parallel reaction monitoring (PRM) acquisition parameters were the following: resolution 17,500; collision energies were set individually in HCD (high-energy collisional dissociation) mode. Metabolites were identified and quantified by accurate mass and retention time and by comparison to the retention times, mass spectra, and responses of known amounts of authentic standards using TraceFinder 4.1 EFS software (ThermoFisher). Retention time alignment, peak picking, deconvolution and normalization by the total ion current (TIC) was performed on Progenesis QI software (version 3.0.6039.34628; Waters). Statistical analysis was performed on SIMCA v. 17.0, MetaboAnalyst v. 6.0 and GraphPad Prism v. 10.6.0.

#### LC-MS/MS

Data acquisition was performed using an adaptation of a method previously described^46^. Samples were injected into using an Dionex UltiMate 3000 LC system (Thermo Scientific) with a Fortis C18 (100 x 2.1 mm; 3 µm) column (Fortis Technology LTD). Solvent A was 0.1 % of acetic acid in water (Optima HPLC grade, Sigma Aldrich) and solvent B was Acetonitrile (Optima HPLC grade, Sigma Aldrich). The column temperature and flow rate were set as 50°C and 0.2 mL min^-1^, and the injection volume was 5 µL. The gradient elution was as follows: 0 % B hold for 3 min, 0 – 60 % B from 3 to 5 min, hold for 2 min, then in 7.01 return to the initial condition and hold for 4 min to re-equilibrate the analytical column. MS was performed in positive and negative polarities using an TQS Quantiva Triple Quadrupole Mass Spectromer (Thermo Scientific) with an electrospray ionization (ESI) source. Qualitative and quantitative analysis was performed using Xcalibur Qual Browser and Tracefinder 5.1 software (Thermo Scientific) according to the manufacturer’s workflows. Analyses were performed in selected reaction monitoring (SRM). Precursor to product ions transitions and collision energies are listed in Table 1.

**Table 1.**
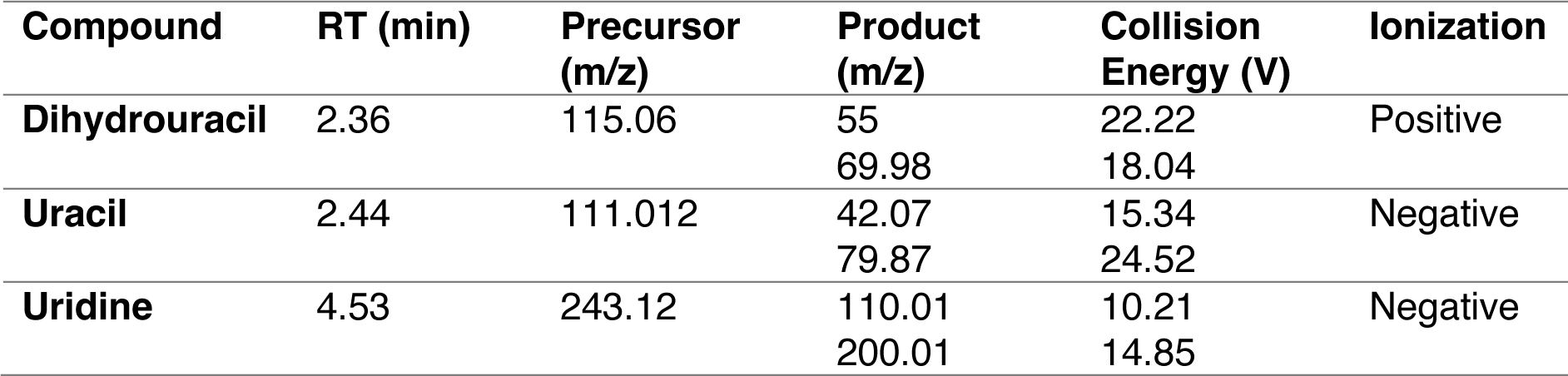
SRM parameters.

#### Chemical and material

Optima water (Optima HPLC grade, Sigma Aldrich)

Acetonitrile (Optima HPLC grade, Sigma Aldrich)

Acetic acid (Fisher Chemical)

Chloroform (Optima HPLC grade, Sigma Aldrich)

Methanol (Optima HPLC grade, Sigma Aldrich)

Ammonium carbonate (Sigma Aldrich, SKU 207861)

13C5, ^15^N_1_-valine (Cambridge Isotope Laboratories, Item number CNLM-442-H-0.25)

Human serum (Sigma Aldrich, H4522)

Dihydrouracil (Sigma Aldrich, 504-07-4)

Uracil (Sigma Aldrich, 66-22-8)

Uridine (Sigma Aldrich, 58-96-8)

## ACKNOWLEDGMENTS

We are grateful to Chris Tape (UCL, London) for advice with scRNA analysis, and Andras Wack and Cecilia Johansson for advice on lung infections. We thank Anna Bualies Domenech for assistance with intestinal organoid assays. We thank the core facilities at the Francis Crick Institute that assisted with this work, including the Biological Research Facility (Tim Zverev, Jack Williams, Nicolas Chisholm and Ola Puchalska-Osorio), Metabolomic, Flow Cytometry Facility (Muhammad Saeed, Ana Agua-Doce, Sina Nanjou and Steve Lim), Experimental Histopathology Laboratory (Richard Stone and Emma Nye), Light Microscopy (Matt Renshaw and Donald Bell), Advanced Sequencing Facility (María Rodríguez, Olga O’Neill, Hubert Slawinski, Robert Gunn) and Bioinformatic (Lina Gerontogianni, James Cambell). Schematics were created with BioRender. This work was supported by the Francis Crick Institute, which receives its core funding from Cancer Research UK (CC2051), the UK Medical Research Council (CC2051), the Wellcome Trust (CC2051), and the European Research Council grant (ERC CoG-H2020725492).

## AUTHOR CONTRIBUTIONS

N.R. designed and performed experiments, analysed most of the data, interpreted the results and co-wrote the manuscript. V.L.B. managed colony breeding and performed some of the experiments; C.P., F.R., A.F, R.M.M.F., K.A., O.N. contributed to some experiments and provided technical support, P.C., L.G., performed some of the bioinformatic analysis (RNA-seq); M.S. performed the blood imaging flow cytometry; M.H. N.L., M.H, I.F, N.L and J.M. performed the metabolomic analysis and help design the protocols. I.M. designed and supervised the study, interpreted the data, and co-wrote the manuscript.

## References

1. Rabas, N., Ferreira, R. M. M., Blasio, S. D. & Malanchi, I. Cancer-induced systemic pre-conditioning of distant organs: building a niche for metastatic cells. Nat. Rev. Cancer 24, 829– 849 (2024).

2. Patras, L., Shaashua, L., Matei, I. & Lyden, D. Immune determinants of the pre-metastatic niche. Cancer Cell 41, 546–572 (2023).

3. Kaplan, R. N. et al. VEGFR1-positive haematopoietic bone marrow progenitors initiate the pre-metastatic niche. Nature 438, 820–827 (2005).

4. Hongu, T. et al. Perivascular tenascin C triggers sequential activation of macrophages and endothelial cells to generate a pro-metastatic vascular niche in the lungs. *Nat*. Cancer 3, 486– 504 (2022).

5. Erler, J. T. et al. Hypoxia-Induced Lysyl Oxidase Is a Critical Mediator of Bone Marrow Cell Recruitment to Form the Premetastatic Niche. Cancer Cell 15, 35–44 (2009).

6. Midwood, K. S., Mao, Y., Hsia, H. C., Valenick, L. V. & Schwarzbauer, J. E. Modulation of Cell–Fibronectin Matrix Interactions during Tissue Repair. J. Investig. Dermatol. Symp. Proc. 11, 73–78 (2006).

7. Hedrick, C. C. & Malanchi, I. Neutrophils in cancer: heterogeneous and multifaceted. Nat. Rev. Immunol. 22, 173–187 (2021).

8. Nolan, E. & Malanchi, I. Connecting the dots: Neutrophils at the interface of tissue regeneration and cancer. Semin Immunol 57, 101598 (2022).

9. Deyell, M., Garris, C. S. & Laughney, A. M. Cancer metastasis as a non-healing wound. Br. J. Cancer 124, 1491–1502 (2021).

10. Reinecke, J. B. et al. Aberrant Activation of Wound-Healing Programs within the Metastatic Niche Facilitates Lung Colonization by Osteosarcoma Cells. Clin. Cancer Res. 31, 414–429 (2025).

11. Ombrato, L. et al. Metastatic-niche labelling reveals parenchymal cells with stem features. Nature 572, 603–608 (2018).

12. Ombrato, L. et al. Generation of neighbor-labeling cells to study intercellular interactions in vivo. Nature Protocols 16, 872–892 (2021).

13. Rodrigues, F. S. et al. Bidirectional activation of stem-like programs between metastatic cancer and alveolar type 2 cells within the niche. Dev. Cell 59, 2398–2413.e8 (2024).

14. Wculek, S. K. & Malanchi, I. Neutrophils support lung colonization of metastasis-initiating breast cancer cells. Nature 528, 413–7 (2015).

15. Coffelt, S. B. et al. IL-17-producing γδ T cells and neutrophils conspire to promote breast cancer metastasis. Nature 522, 345–348 (2015).

16. Nolan, E. et al. Radiation exposure elicits a neutrophil-driven response in healthy lung tissue that enhances metastatic colonization. *Nat*. Cancer 3, 173–187 (2022).

17. Liu, K. et al. Tracing the origin of alveolar stem cells in lung repair and regeneration. Cell 187, 2428–2445.e20 (2024).

18. Moon, K. R. et al. Visualizing structure and transitions in high-dimensional biological data. Nat. Biotechnol. 37, 1482–1492 (2019).

19. Choi, J. et al. Inflammatory Signals Induce AT2 Cell-Derived Damage-Associated Transient Progenitors that Mediate Alveolar Regeneration. Cell Stem Cell 27, 366–382.e7 (2020).

20. Strunz, M. et al. Alveolar regeneration through a Krt8+ transitional stem cell state that persists in human lung fibrosis. Nat. Commun. 11, 3559 (2020).

21. Rawlins, E. L. Lung Epithelial Progenitor Cells. Proc. Am. Thorac. Soc. 5, 675–681 (2008).

22. Burkhardt, D. B. et al. Quantifying the effect of experimental perturbations at single-cell resolution. Nat. Biotechnol. 39, 619–629 (2021).

23. Garner, H. et al. Understanding and reversing mammary tumor-driven reprogramming of myelopoiesis to reduce metastatic spread. Cancer Cell (2025) doi:10.1016/j.ccell.2025.04.007.

24. LaMarche, N. M. et al. An IL-4 signalling axis in bone marrow drives pro-tumorigenic myelopoiesis. Nature 625, 166–174 (2024).

25. Ramessur, A. et al. Circulating neutrophils from patients with early breast cancer have distinct subtype-dependent phenotypes. Breast Cancer Res. 25, 125 (2023).

26. Whyte, D. et al. Uridine phosphorylase-1 supports metastasis by altering immune and extracellular matrix landscapes. EMBO Rep. 26, 4248–4282 (2025).

27. Manz, M. G. & Boettcher, S. Emergency granulopoiesis. Nat. Rev. Immunol. 14, 302–314 (2014).

28. Xie, X. et al. Single-cell transcriptome profiling reveals neutrophil heterogeneity in homeostasis and infection. Nat Immunol 21, 1–37 (2020).

29. Cao, D., Leffert, J. J., McCabe, J., Kim, B. & Pizzorno, G. Abnormalities in Uridine Homeostatic Regulation and Pyrimidine Nucleotide Metabolism as a Consequence of the Deletion of the Uridine Phosphorylase Gene*. J. Biol. Chem. 280, 21169–21175 (2005).

30. Browaeys, R., Saelens, W. & Saeys, Y. NicheNet: modeling intercellular communication by linking ligands to target genes. Nat. Methods 17, 159–162 (2020).

31. Lecot, P. et al. Gene signature of circulating platelet-bound neutrophils is associated with poor prognosis in cancer patients. Int. J. Cancer 151, 138–152 (2022).

32. Wang, J. et al. Visualizing the function and fate of neutrophils in sterile injury and repair. Science 358, 111–116 (2017).

33. Phillipson, M. & Kubes, P. The Healing Power of Neutrophils. Trends in Immunology 40, 635–647 (2019).

34. Sreeramkumar, V. et al. Neutrophils scan for activated platelets to initiate inflammation. Science 346, 1234–1238 (2014).

35. Rafii, S. et al. Platelet-derived SDF-1 primes the pulmonary capillary vascular niche to drive lung alveolar regeneration. Nat. Cell Biol. 17, 123–136 (2015).

36. Labelle, M., Begum, S. & Hynes, R. O. Direct Signaling between Platelets and Cancer Cells Induces an Epithelial-Mesenchymal-Like Transition and Promotes Metastasis. Cancer Cell 20, 576–590 (2011).

37. Schumacher, D., Strilic, B., Sivaraj, K. K., Wettschureck, N. & Offermanns, S. Platelet-Derived Nucleotides Promote Tumor-Cell Transendothelial Migration and Metastasis via P2Y2 Receptor. Cancer Cell 24, 130–137 (2013).

38. Lucotti, S. et al. Aspirin blocks formation of metastatic intravascular niches by inhibiting platelet-derived COX-1/thromboxane A2. J. Clin. Invest. 130, 1845–1862 (2019).

39. Hergueta-Redondo, M. et al. The impact of a high fat diet and platelet activation on pre-metastatic niche formation. Nat. Commun. 16, 2897 (2025).

40. Malanchi, I. et al. Interactions between cancer stem cells and their niche govern metastatic colonization. Nature 481, 85–89 (2011).

41. Gracz, A. D., Puthoff, B. J. & Magness, S. T. Somatic Stem Cells, Methods and Protocols. Methods Mol. Biol. 879, 89–107 (2012).

42. Jose, L. H. S. & Signer, R. A. J. Cell-type-specific quantification of protein synthesis in vivo. Nat. Protoc. 14, 441–460 (2019).

43. Chen, E. Y. et al. Enrichr: interactive and collaborative HTML5 gene list enrichment analysis tool. BMC Bioinform. 14, 128 (2013).

44. Kuleshov, M. V. et al. Enrichr: a comprehensive gene set enrichment analysis web server 2016 update. Nucleic Acids Res. 44, W90–W97 (2016).

45. Xie, Z. et al. Gene Set Knowledge Discovery with Enrichr. Curr. Protoc. 1, e90 (2021).

46. Fugger, K. et al. Targeting the nucleotide salvage factor DNPH1 sensitizes BRCA-deficient cells to PARP inhibitors. Science 372, 156–165 (2021).

