## Extended Data Figures for "Cancer-driven neutrophil priming couples systemic epithelial regenerative programs with pre-metastatic niche formation"

#### Extended Data Figure 1

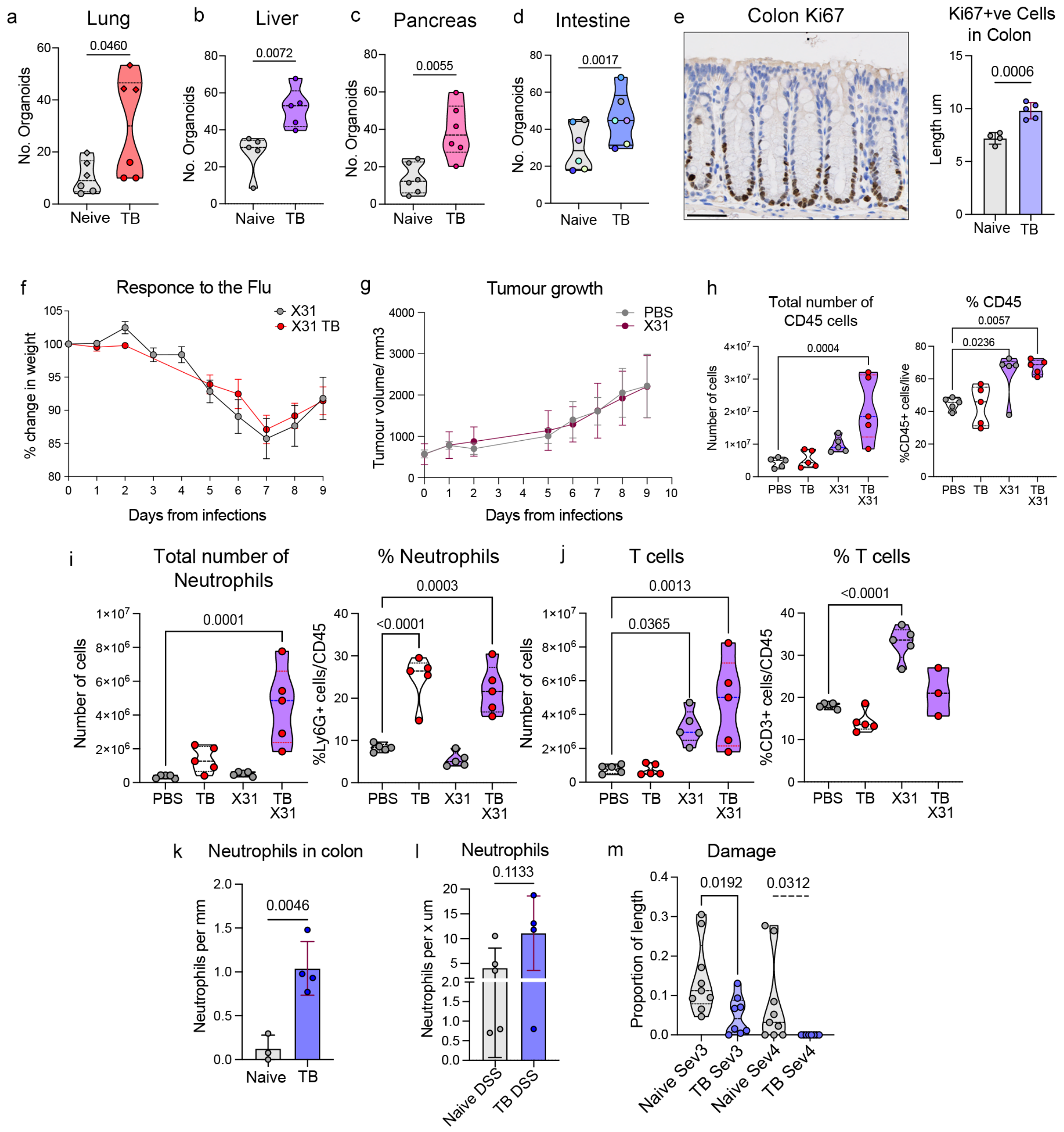


**Extended Data Figure 1 Systemic enhancement of epithelial progenitor activity and tissue responses in tumour-bearing mice.**
**a-d)** Total organoid number from cells isolated from lung (a), liver (b), pancreas (c) and intestine (d) of tumour-bearing (TB) and naïve mice (PyMT, FVB). Violin plots show organoid numbers per mouse (n= 6 per group except liver n=5). For lung, liver, pancreas, two-tailed Student's t-test was used. Intestinal organoids were analysed with paired t-test. **e) (Left)** Representative image of proliferating cells in colon crypts based Ki67 immunohistochemistry. **(Right)** Histological quantification. Data represent mean number of K67+ cells per crypt (naïve n=4, TB n=5) Welch's t-test. **f)** Body weight monitoring following influenza infection as percentage change from baseline (naïve n=5, TB n=5 PyMT, C57/Bl6). The tumour weight over time was calculated from the tumour volume and subtracted from the wight of TB mice. **g)** Tumour burden following influenza infection represented as tumour volume (naïve n=5, TB n=5). **h-j)** Quantification of **(left)** total or **(right)** proportion of immune cell populations in lungs of naïve and TB mice: CD45+ cells (h), Neutrophils, (i) and T-cells (j) (naïve n=5, TB n=5). Ordinary one-way ANOVA. **k–l)** Quantification of neutrophils in the colon of naïve and TB mice (PyMT, FVB) (k), and following DSS treatment (l), based on S100A9 immunohistochemistry.

Data represent the mean number of neutrophils per mm of colon (naïve n=3/5, TB n=4), Welch's t-test. **m)** Direct comparison of proportion of colon length exhibiting damage severity score 3 and 4 in neive and TB mice (data shown in Figure 1j) (naïve n=9, TB n=8). Two-tailed Student's t-test.

### Extended Data Figure 2

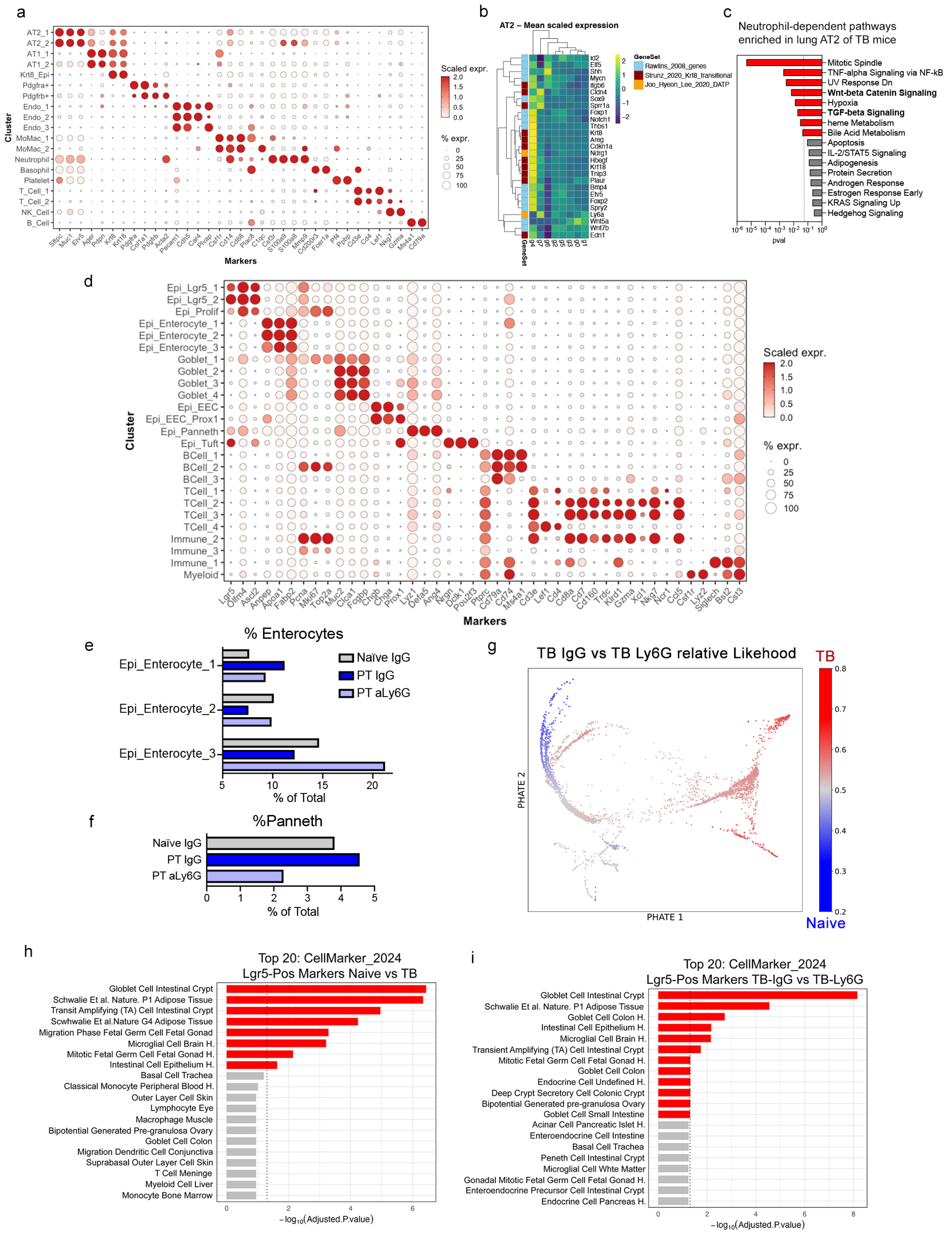

**Extended Data Figure 2 Primary breast tumours induce a neutrophil-dependent perturbation in epithelial compartments of both premetastatic lung and intestine.**
**a)** Canonical marker-gene expression across lung cell-type clusters from Naïve IgG, tumour-bearing (TB) IgG and TB αLy6G, datasets (related to Figure 2a) (PyMT, FVB). Colour scale indicates scaled expression level. **b)** AT2 cluster-wide mean scaled expression level of genes associated with alveolar type 2 (AT2) progenitor-associated signatures (related to Figure 2c).

Individual gene sets and source are shown in the legend. g1-7 correspond to AT2 PHATE clusters 1-7 shown Figure 2c. **c)** Neutrophil-dependent pathways enriched in lung AT2 cells from TB mice. Pathways are derived from MSigDB Hallmarks 2020 library with enrichment p values indicated. **d)** Canonical marker gene expression across intestinal cell type clusters from Naïve IgG, TB IgG and TB $\alpha$Ly6G dataset related to Figure 2f. Colour represents scaled expression level. **e-f)** Proportions of Enterocyte (e) and Paneth cells (f) proportions within intestinal scRNAseq data calculated for Naïve IgG, TB IgG and TB $\alpha$Ly6G conditions. **g)** Intestinal epithelial per-cell MELD relative likelihood represented on PHATE embedding for TB IgG and TB $\alpha$Ly6G conditions. Red is associated with higher relative likelihood associated with TB IgG condition. **h-i)** Pathway enrichment for the Lgr5-positive intestinal epithelial cluster comparing TB IgG vs Naïve IgG conditions (h) or TB IgG and TB $\alpha$Ly6G (i) conditions.

#### Extended Data Figure 3

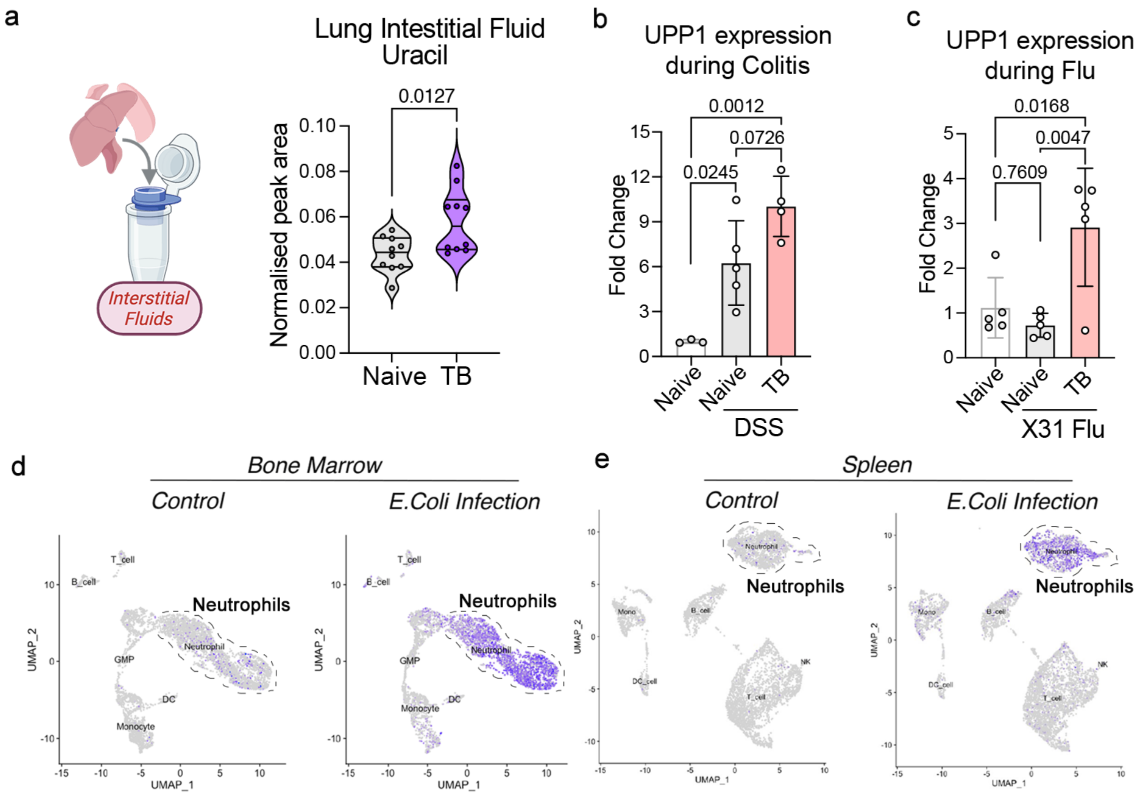


**Extended Data Figure 3. Cance-primed neutrophils acquire UPP1 expression.**
**a) (Left)** Schematic for LCMS-based metabolic analysis of lung interstitial fluid (LIF). Freshly isolated lungs were dried processed by differential centrifugation over a filter to collect LIF for LC–MS analysis. **(Right)** Uracil abundance in LIF from Naïve and TB mice (4T1, Balb/c) (n = 10), Welch's t-test. **b)** UPP1 mRNA expression determined by qPCR in bone-marrow neutrophils isolated from naïve control mice, or from naïve and TB (PyMT, FVB) mice treated with DSS. Data represent fold change in expression relative to untreated naïve controls (n=3 naïve, n=5 naïve + DSS, n=4 TB + DSS); ordinary one-way ANOVA. **c)** UPP1 mRNA expression determined by qPCR in bone marrow neutrophils isolated from naïve control mice or influenza infected naïve or TB mice (PyMT, C57/Bl6) at day 9 post infection (n=5 per group), ordinary one-way ANOVA. **d-e)** UPP1 expression from published scRNAseq datasets of bone marrow and splenic cells isolate from control or E.coli-infected mice (Xie, X. et al.2020)[28]. UMAPs indicate UPP1 expression intensity (purple) across annotated cell types, dotted line highlights Neutrophil pool.

### Extended Data Figure 4

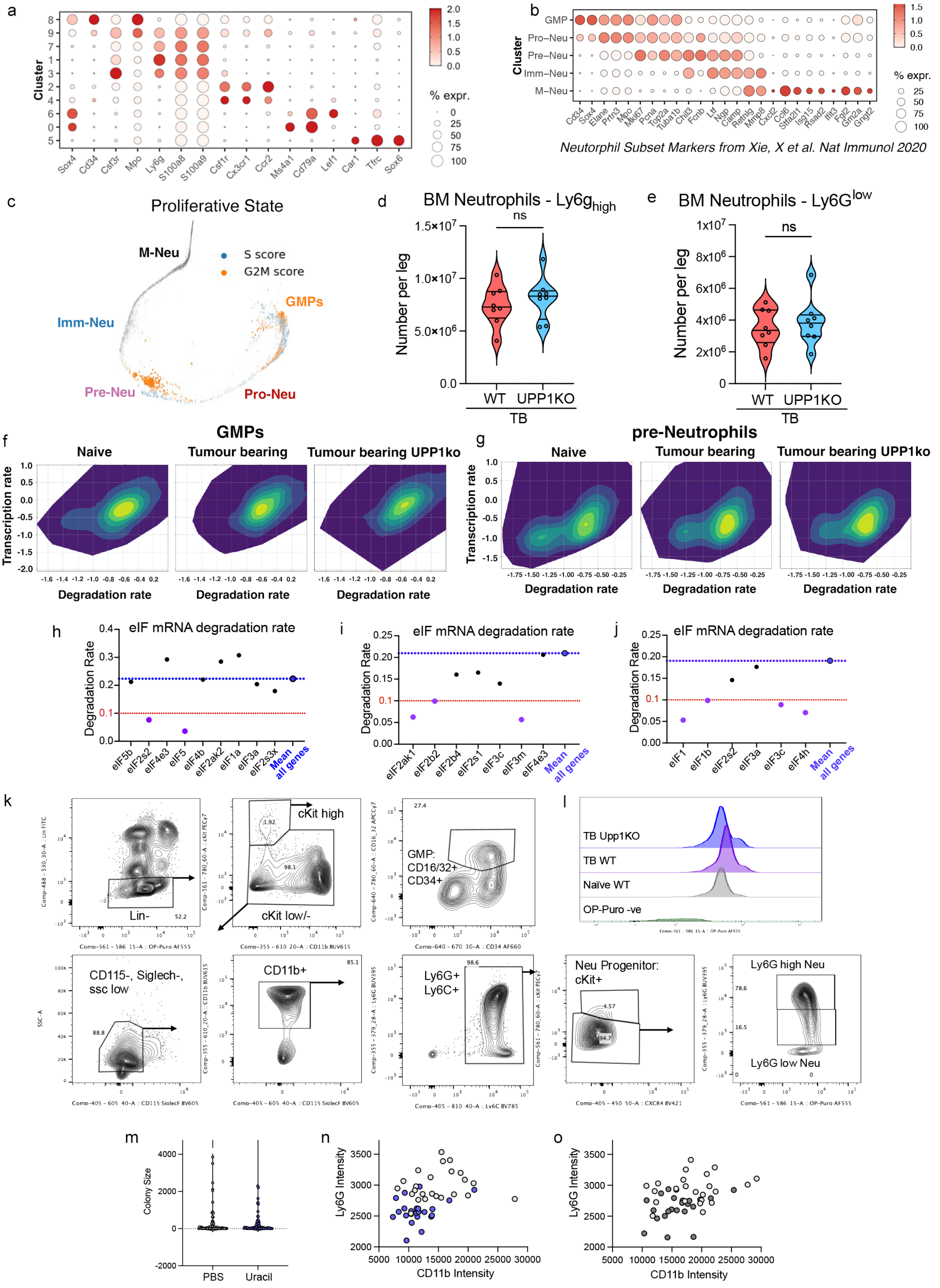

**Extended Data Figure 4. Characterisation of granulopoiesis in UPP1KO mice.**
**a)** Canonical marker-gene expression used to identify broad cell types across bone marrow (BM) clusters from Naïve WT tumour bearing (TB) WT and TB UPP1KO samples (PyMT, C57/Bl6) (related to Figure 4a). Colour scale indicates scaled expression level. **b)** Marker gene expression used to identify neutrophil and neutrophil progenitor subtypes within granulocytic cell lineage related to Figure 4b. **c)** Cell cycle state prediction across BM neutrophils and neutrophil progenitors. **d-e)** Absolute number of bone marrow mature (d) and immature (e) neutrophils in TB WT and TB UPP1KO mice (n=8 per group) Welch's t-test. **f-g)** mRNA dynamics across neutrophil and progenitor populations**.** Predicted transcription (y-axis) and degradation (x-axis) rates for individual genes in GMP (f) and pre-Neutrophil (g) clusters from Naïve WT, TB WT and TB Upp1KO mice, derived using scVelo dynamical modelling. **h-j)** Predicted degradation rates of eukaryotic translation initiation factor (eIF) transcripts within the GMP (h), pre-neutrophil (i) and immature neutrophil (j) populations. Mean degradation rate for all transcripts is indicated in blue; 'protected' transcripts with degradation rate < 0.1 are shown in purple. **k)** Representative flow cytometry plots showing the gating strategy used to identify GMP, pre/pro neutrophils, immature and mature neutrophils for OP-Puro fluorescent analysis. **l)** Example of plot showing the fluorescent signal from OP-Puro. **m)** Monocyte ($Ly6C^+$) cell number per colony from GMP colony-formation assay with or without uracil (n=6 mice. 62 PBS and 66 uracil-treated colonies) Mann-Whitney test. **n-o)** Scatter-plot representation of data from Figure 4p. Each point represents mean CD11b and Ly6G fluorescence intensity for circulating WT (light grey) or UPP1KO (blue) $CD45.2^+$ donor neutrophils (n) and for WT $CD45.1^+$ recipient neutrophils from chimeric mice reconstituted with WT (light grey) or UPP1KO (dark grey) CD45.2 bone marrow (o). (WT n=24, UPP1KO n=23.)

#### Extended Data Figure 5

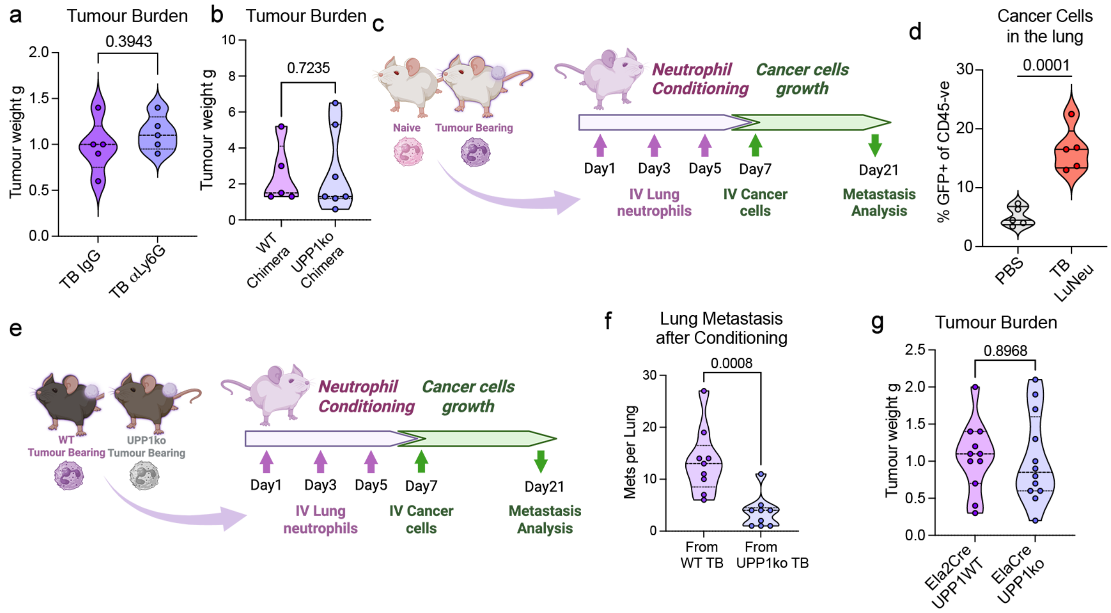


**Extended Data Figure 5. UPP1-deficient cancer primed neutrophils show reduced lungs pre-metastatic activity.**

**a–b)** Tumour burden corresponding in mice from Figures 5a (PyMT, FVB) and 5e (PyMT, C57/Bl6), respectively, (a) n=5 per group, (b) WT Chimera n=5, UPP1KO Chimera n=7. **c)** Schematic of experimental approach to assess the pro-metastatic conditioning potential of neutrophils. Lung neutrophils were isolated by magnetic sorting (MACS) from tumour-bearing mice (PyMT, FVB) and intravenously (IV) transferred into naïve recipients. Three rounds of neutrophil transfer were performed over 7 days before IV injection of primary $GFP^{+}$ MMTV-PyMT tumour cells. **d)** Metastatic burden in neutrophil-conditioned mice (from (c)), quantified as the percentage of $GFP^{+}$ PyMT tumour cells (FVB) among $CD45^{-}$ lung cells (n=5 per group), unpaired t-test. **e)** Schematic of experimental approach to assess UPP1-dependent pro-metastatic conditioning. Neutrophils were isolated from tumour-bearing WT or UPP1KO mice (PyMT, C57/Bl6) and IV injected into naïve recipients. Three transfers were performed before IV injection of primary MMTV-PyMT tumour cells. **f)** Quantification of lung metastasis based on histological sections from mice conditioned with neutrophils from tumour-bearing WT or UPP1KO donor mice. Data represent number of metastatic foci per lung. (n=9 mice per group) unpaired t-test. **g)** Tumour burden related to figure 5h (WT n=11, Ela2CRE Upp1KO n=12)

#### Extended Data Figure 6

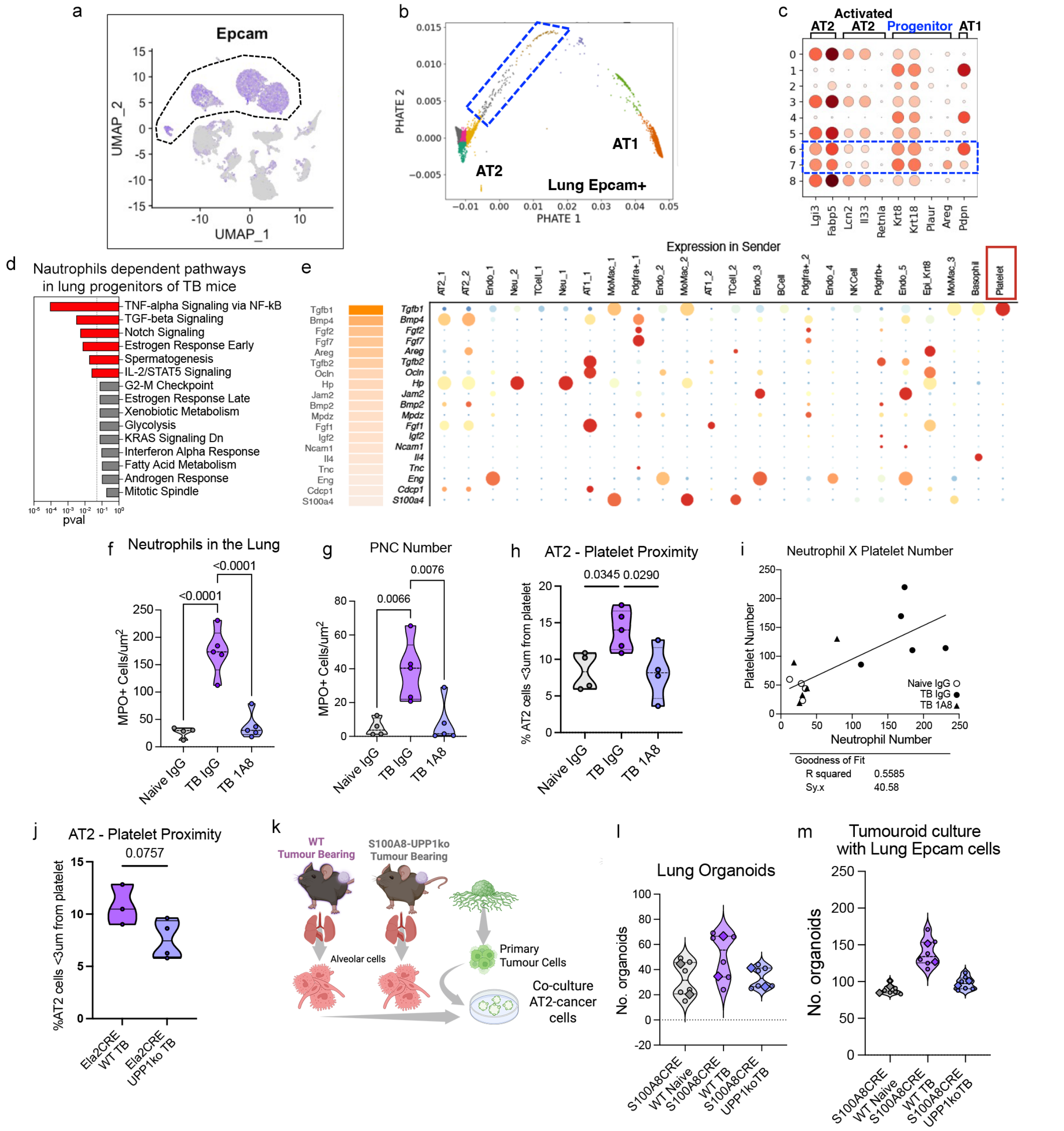


**Extended Data Figure 6. Transcriptionally altered AT2 are predicted to interact with platelets.**
**a-d)** Analysis workflow assessing neutrophil-dependent transcriptional programs in epithelial progenitors within the premetastatic lung (PyMT, FVB). (b) Epcam$^+$ epithelial cells from the premetastatic lung scRNA-seq dataset (UMAP shows Epcam expression in purple) were used to (c) generate a PHATE embedding capturing epithelial lineage hierarchy. (d) Canonical marker-gene expression (DotPlot) was used to identify alveolar epithelial progenitor clusters, followed by (e) pathway enrichment analysis comparing TB IgG and TB αLy6G conditions. **e)** Ligand-receptor analysis using NicheNet to identify microenvironmental interactions driving the neutrophil-dependent transcriptional perturbation of alveolar epithelial progenitors identified in (b). **(Left)** Ranked ligand activity scores quantifying the ability of each ligand to explain the TB IgG versus TB αLy6Gg transcriptional changes. **(Right)** expression of the top-ranked ligands across all cell-type clusters within the premetastatic lung, showing that platelets exhibited the highest expression of the top-ranked ligand, Tgfβ1. **f)** Quantification of neutrophil

number in perfused lung of naïve IgG, TB IgG and TB αLy6G mice (PyMT, FVB). Data represent number of MPO+ neutrophils $mm^2$ lung tissue (naïve IgG n=4, TB IgG and TB αLy6G n=5), ordinary one-way ANOVA. **g)** Quantification of platelet-neutrophil cluster (PNC) number in perfused lung of naïve IgG, TB IgG and TB αLy6G mice (PyMT, FVB). Data represent number of MPO+ neutrophils in close contact with CD41b+ platelets per $mm^2$ lung tissue (naïve IgG n=4, TB IgG and TB αLy6G n=5), ordinary one-way ANOVA. **h)** Quantification of AT2 cells ($SPC^+$) in close proximity (<3 μm) to platelets ($CD41b^+$) in perfused lungs of naïve IgG, TB IgG and TB αLy6G mice (PyMT, FVB) (naïve IgG n=4, TB IgG and TB αLy6G n=5), ordinary one-way ANOVA. **i)** Correlation between neutrophil and platelet numbers within perfused lung of naïve IgG (open circles), TB IgG (filled circles) and TB αLy6G (filled triangles) mice (PyMT, FVB). Neutrophil number plotted on the x-axis and platelet number on the y-axis; simple linear regression shown. **j)** Quantification of AT2 cells (SPC+) in close contact with platelets (CD41b+) in perfused lung of TB Ela2CRE-WT and TB Ela2CRE-UPP1KO mice (PyMT, C57/Bl6). Data represent percentage of AT2 cells < 3 um from platelet (TB Ela2CRE-WT n=3 and TB Ela2CRE-UPP1KO n=4) unpaired t-test. **k)** Schematic of experimental design to assess lung epithelial organoid formation in mice with neutrophil specific UPP1KO TB mice. Primary MMTV-PyMT tumour cells (C57/Bl6). were co-cultured with lung epithelial cells isolated from naïve S100A8CRE-WT, TB S100A8CRE-WT and TB S100A8CRE-UPP1KO mice (PyMT, C57/Bl6). **l)** Lung epithelial organoid formation efficiency from naïve S100A8Cre-WT, TB S100A8CRE-WT and TB S100A8CRE-UPP1KO mice (PyMT, C57/Bl6) (n=2 biological replicates per group, technical replicates shown in small points). **m)** Tumour organoid formation efficiency in co-cultured with lung epithelial cells isolated form naïve S100A8CRE-WT, TB S100A8CRE-WT and TB S100A8CRE-UPP1KO mice (PyMT, C57/Bl6) (n=2 biological replicates per group with technical replicates shown in small datapoints).
